# On the possibility of yet a third kinetochore system in the protist phylum Euglenozoa

**DOI:** 10.1101/2024.08.06.606595

**Authors:** Corinna Benz, Maximilian W. D. Raas, Pragya Tripathi, Drahomíra Faktorová, Eelco C. Tromer, Bungo Akiyoshi, Julius Lukeš

**Author notes:** Corinna Benz, Maximilian W. D. Raas, and Pragya Tripathi contributed equally to this work. Address correspondence to Julius Lukeš, Bungo Akiyoshi or Eelco C. Tromer.

## Abstract

Transmission of genetic material from one generation to the next is a fundamental feature of all living cells. In eukaryotes, a macromolecular complex called the kinetochore plays crucial roles during chromosome segregation by linking chromosomes to spindle microtubules. Little is known about this process in evolutionarily diverse protists. Within the supergroup Discoba, Euglenozoa forms a speciose group of unicellular flagellates - kinetoplastids, euglenids, and diplonemids. Kinetoplastids have an unconventional kinetochore system, while euglenids have subunits that are conserved amongst most eukaryotes. For diplonemids, a group of extremely diverse and abundant marine flagellates, it remains unclear what kind of kinetochores are present. Here, we employed deep homology detection protocols using profile-versus-profile Hidden Markov Model searches and AlphaFold-based structural comparisons to detect homologies that might have been previously missed. Interestingly, we still could not detect orthologs for most of the kinetoplastid nor canonical kinetochore subunits with few exceptions including a putative centromere-specific histone H3 variant (cenH3/CENP-A), the spindle checkpoint protein Mad2, the chromosomal passenger complex members Aurora and INCENP, and broadly conserved proteins like CLK kinase and the meiotic synaptonemal complex proteins SYCP2/3 that also function at kinetoplastid kinetochores. We examined the localization of five candidate kinetochore-associated proteins in the model diplonemid, *Paradiplonema papillatum*. *Pp*CENP-A shows discrete dots in the nucleus, implying that it is likely a kinetochore component. *Pp*Mad2, *Pp*CLK^KKT10/19^, *Pp*SYCP2L1^KKT17/18^ and INCENP reside in the nucleus, but no clear kinetochore localization was observed. Altogether, these results point to the possibility that diplonemids evolved a hitherto unknown type of kinetochore system.

**IMPORTANCE:** A macromolecular assembly called the kinetochore is essential for the segregation of genetic material during eukaryotic cell division. Therefore, characterization of kinetochores across species is essential for understanding the mechanisms involved in this key process across the eukaryotic tree of life. In particular, little is known about kinetochores in divergent protists such as Euglenozoa, a group of unicellular flagellates that includes kinetoplastids, euglenids, and diplonemids, the latter being a highly diverse and abundant component of marine plankton. While kinetoplastids have an unconventional kinetochore system and euglenids have a canonical one similar to traditional model eukaryotes, preliminary searches detected neither unconventional nor canonical kinetochore components in diplonemids. Here, we employed state-of-the-art deep homology detection protocols but still could not detect orthologs for the bulk of kinetoplastid-specific nor canonical kinetochore proteins in diplonemids except for a putative centromere-specific histone H3 variant. Our results strongly suggest that diplonemids evolved kinetochores that do not resemble previously known ones.

---

A fundamental characteristic of living organisms is the ability to self-replicate. This ability requires accurate transmission of genetic information from one generation to the next during cell division (1, 2). In eukaryotes, a sophisticated macromolecular structure called the kinetochore assembles on centromeric chromatin and binds spindle microtubules to control chromosome movement during mitosis and meiosis (3). In addition, kinetochores are responsible for regulating a surveillance mechanism called the spindle assembly checkpoint (SAC), which delays cell cycle progression until all chromosomes have achieved proper kinetochore-microtubule attachment (4). The molecular mechanisms underlying these processes have been extensively investigated in traditional model eukaryotes, such as yeast, worm, fly, and human. Importantly, all of these organisms belong to the supergroup Opisthokonta, meaning that they are closely related in the timescale of eukaryotic evolution (**Fig. 1A**) (5). With few exceptions, little is known about mitotic mechanisms in members of other eukaryotic supergroups (6).

**Fig 1.**
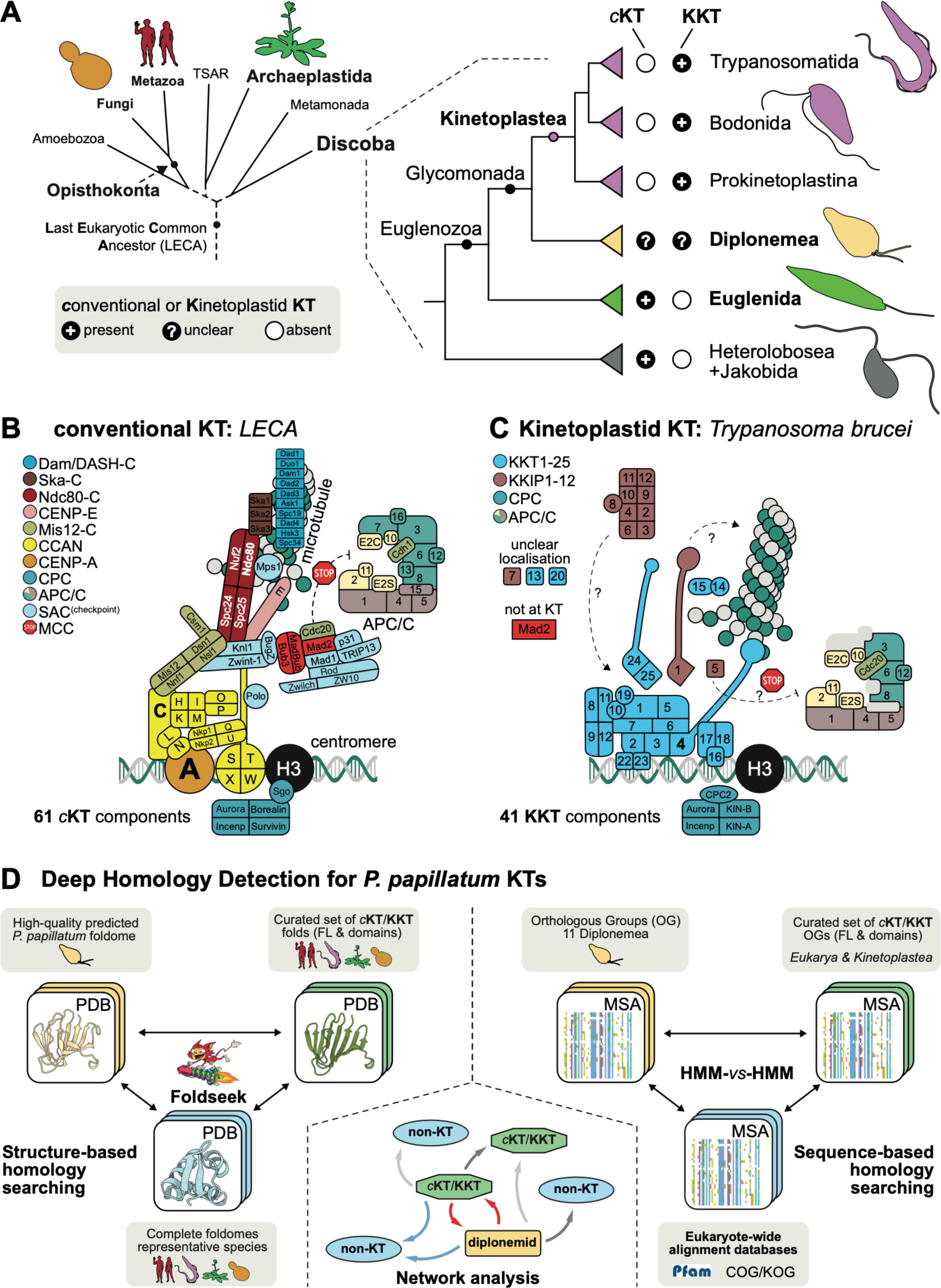
Kinetochore composition in Euglenozoa. (**A**) Cartoon of phyletic relationships of the superphylum Discoba with emphasis on the phylum Euglenozoa harboring kinetoplastids, euglenids, and diplonemids. (**B**) Cartoon of 61 components of the *conventional* KT (*c*KT), spindle assembly checkpoint (SAC), and the anaphase promoting complex/cyclosome (APC/C) as inferred to have been present in the ancestral eukaryotes (11). (**C**) Cartoon of 40 components and structure of the kinetoplastid kinetochore (KKT) in *T. brucei*, also including the SAC and APC/C components. (**D**) Overview of the twofold strategy taken towards highly-sensitive identification of *c*KT/KKT orthologs in the *P. papillatum* proteome. Schematic overviews of the reciprocal structure-based (left) and sequence-based homology searching strategies (right). In the center, an example of a network graph is shown that is produced based on these comprehensive searches, with green octagonal nodes indicating a *c*KT/KKT model, a rectangular yellow node a *P. papillatum* accession or Diplonemid Orthologous Group (OG), and in circular blue nodes any non-KT protein from the complete foldomes of any of the representative species (*Homo sapiens, Saccharomyces cerevisiae, Arabidopsis thaliana, Trypanosoma brucei*). Arrows (edges) between nodes represent homology links colored according to hierarchy in the search results based on E-value (red: best, blue: second-best, dark grey: third-best and light grey: other hit).

Studies in opisthokonts have shown that the kinetochore is a highly complex structure (more than 100 proteins in humans), which can be divided into two functional modules: the inner kinetochore that is built on centromeres and the outer kinetochore that binds microtubules and regulates the SAC (7, 8). A key inner kinetochore component is CENP-A, a centromere-specific histone H3 variant that determines the position of kinetochores and initiates kinetochore assembly, while the outer kinetochore includes the Ndc80 complex that binds microtubules. The fact that CENP-A, its interaction partner CENP-C, the Ndc80 complex, and SAC proteins such as Mad2 and Cdc20 are widely conserved suggests that most eukaryotes use these proteins to perform basic kinetochore functions. However, phylogenetic profiling surveys across eukaryotes revealed remarkable plasticity of kinetochore composition (6, 9, 10). Strikingly, reconstructions based on these disparate presence/absence profiles of kinetochore proteins still suggest that the Last Eukaryotic Common Ancestor (LECA) possessed a highly complex ancestral kinetochore system on par with those of extant eukaryotes such as human and yeast (**Fig. 1B**) (11, 12). Such intuitively opposing patterns of ancestral complexity and contemporary plasticity imply that the kinetochores must have had a highly dynamic evolutionary history, despite their cardinal role in chromosome segregation.

Recent experimental studies started to probe kinetochore variations among eukaryotes (13). For example, several insects (e.g. butterflies and moths) and early-diverging fungi (e.g. *Mucor circinelloides*) lack CENP-A, although these organisms have retained the Ndc80 complex (14–16). Furthermore, an increasing number of non-traditional model eukaryotes are being studied for their seemingly divergent kinetochore composition and function, such as the malaria parasite *Plasmodium* (17, 18), while an extensive bioinformatic survey could not detect most of the conventional kinetochore in the free-living metamonad flagellate *Carpediemonas membranifera* (19).

The most extreme case of kinetochore divergence known to date is found in Kinetoplastea (Euglenozoa, Discoba) for which bioinformatic analysis failed to identify obvious orthologs of CENP-A, Ndc80, or any other canonical kinetochore proteins (20, 21). A localization-based screen in the model kinetoplastid parasite *Trypanosoma brucei* identified kinetoplastid kinetochore protein 1 (KKT1), and subsequent immunoprecipitation and mass spectrometry analyses identified many additional kinetochore proteins including KKT1–25 and KKIP1–12 (**Fig. 1C**) (22–26). Many of these proteins are conserved among kinetoplastids, including free-living bodonids and prokinetoplastids (**Fig. 1A**) (20, 27–30). However, clear homologs for most KKT/KKIPs are apparently absent in other eukaryotes, except for proteins with either generic domains for which pinpointing their evolutionary history is technically challenging (e.g. PH and FHA domains in KKT proteins and RRM domains in many KKIP proteins), or for proteins such as KKT16/17/18 and KKT10/19 that also play key roles in meiotic chromosome synapsis and splicing in other eukaryotes, respectively. Although it has been proposed that the ancestor of kinetoplastids repurposed parts of the meiotic synapsis and homologous recombination machinery to assemble unique kinetochores (31), it remains unclear why and when kinetoplastids invented their unique kinetochore system.

Kinetoplastids belong to the phylum Euglenozoa, which also includes euglenids and diplonemids (**Fig. 1A**) (32, 33). Phylogenetic studies indicate that kinetoplastids are more closely related to diplonemids than to euglenids (29, 34). Diplonemids are a group of heterotrophic flagellates that are not only highly abundant in marine environments (35–37), but also extremely diverse, with 18S rRNA-based estimates reaching ∼70,000 species (38). Negligence of these omnipresent eukaryotes is reflected by the fact that the number of diplonemid species in culture for which morphological and sequence data is available still does not exceed a dozen (37, 39). Hence, despite an inevitable ecological importance of these highly-abundant protists in the oceans, very little is known about their biology. Interestingly, unlike euglenids that have clear orthologs of canonical kinetochore proteins, only a putative CENP-A candidate has been identified in diplonemids (29). Hence, the kinetochore composition in diplonemids remains unclear.

To address this knowledge gap and to gain insights into the evolutionary origin of kinetoplastid kinetochores, we set out to investigate the possible protein composition of kinetochores in diplonemids. We started by applying state-of-the-art sensitive homology detection protocols to explore the possibility of previously missed hidden homologies (18, 40). Indeed, in the malaria parasite *Plasmodium berghei*, a comparative analysis using both profile Hidden Markov Model (HMMs) and AlphaFold2-predicted protein structures uncovered the presence of many *bona fide* canonical kinetochore proteins that were previously refractory to discovery even by powerful tools such as iterative HMM searches (18). To achieve the same goal, we have used AlphaFold2 to predict the Foldome for the model diplonemid *Paradiplonema papillatum* (formerly known as *Diplonema papillatum*).

Recently, *P. papillatum* became the first diplonemid (Diplonemea) for which methods of integration of extraneous DNA have been developed (41), thus turning it into a new model marine protist (42). Indeed, insertion of any DNA segment into a chromosome *via* homologous recombination can be achieved by using ∼1.5 kb-long homology arms (43). The application of tagging to identify components of membrane trafficking and the mitochondrial ribosome confirmed experimental tractability of *P. papillatum* (44, 45). With a publicly available high-quality genome (46) and a set of transcriptomes (47), this technique enables functional studies in this group of evolutionarily and ecologically highly relevant protists. In this study, we explore the evolutionary history of euglenozoan kinetochores and examine the localization of five candidate kinetochore-associated proteins in *P. papillatum*.

## RESULTS

### Deep homology detection strategies confirm the general absence of known kinetochore proteins amongst Diplonemea

Previously, it was suggested that Diplonemea may lack several crucial components of the conventional kinetochore (*c*KT) and also of the kinetoplastid kinetochore (KKT), leaving these flagellates seemingly without a known type of kinetochore structure (29). Here, we built on these prior analyses by performing a comprehensive and highly-sensitive deep homology search for components of the *c*KT and KKT, focusing on the predicted proteome from *P. papillatum*. This workflow allows us to scrutinise several mutually non-exclusive evolutionary scenarios for the diplonemid kinetochore, which is either: i/ a highly divergent version of the *c*KT; ii/ a highly divergent version of the KKT; iii/ a mix of both *c*KT/KKT, or iv/ a unique kinetochore of distinct evolutionary origin.

To systematically search the proteome of *P. papillatum* for highly divergent orthologs of *c*KT and KKT components, we used a twofold strategy employing both sequence-based as well as predicted protein structure-based similarity searches (**Figs. 1D, S1, S2**). In the former, we performed exhaustive multilevel profile-versus-profile Hidden Markov Model (HMM) searches specifically exploring ‘grey-zone’/borderline similarities leveraging sensitive, manually curated models of both full-length proteins as well as domains and motifs for 71 *c*KT and 39 KKT-related components (**Tables S3, S4**). For structural comparisons (**Table S5**), we predicted the protein structures of the *P. papillatum* proteome with AlphaFold2 (AF2) using multiple sequence alignments including 10 Diplonemea species for which transcriptomic data are available (**Tables S1, S2**). We followed a similar strategy taken in the profile-versus-profile HMM searches, in this case, using 272 manually-curated *c*KT/KKT protein structure AF2 models from four representative species: *Homo sapiens*, *Saccharomyces cerevisiae*, *Arabidopsis thaliana* and *Trypanosoma brucei* (**Fig. 1D, S2**).

Putative homologs obtained by these searches were analyzed in a network graph-based framework using Markov Clustering (MCL) (**Fig. 2; Methods**), with the underlying reasoning that orthologous sequences and folds tend to form distinct similarity clusters in the MCL network that can be used as a proxy for their underlying phylogeny (40, 48). In parallel, we applied the bidirectional best/better hit principle to uncover strong homology associations (**Tables S3C, S5C**). Using these complementary methods, our analysis readily revealed *bona fide* orthologs of the spindle checkpoint (p31^COMET^, Cdc20, Trip13, Mad2), chromosomal passenger complex (INCENP), inner kinetochore histone-like proteins CENP-S and CENP-X, as well as the previously-reported putative CENP-A ortholog (**Fig. 2**). This approach also allowed the identification of multiple close paralogs, and in some cases the closest paralog, to *c*KT/KKT proteins where multiple duplications of a particular gene family occurred among Euglenozoa (e.g. CLK kinase KKT10/19 and the phosphatase KKIP7) (29). Lastly, we recovered two orthologs of the SYCP2-like proteins KKT17 and KKT18, corroborating previous findings (31).

**Fig 2.**
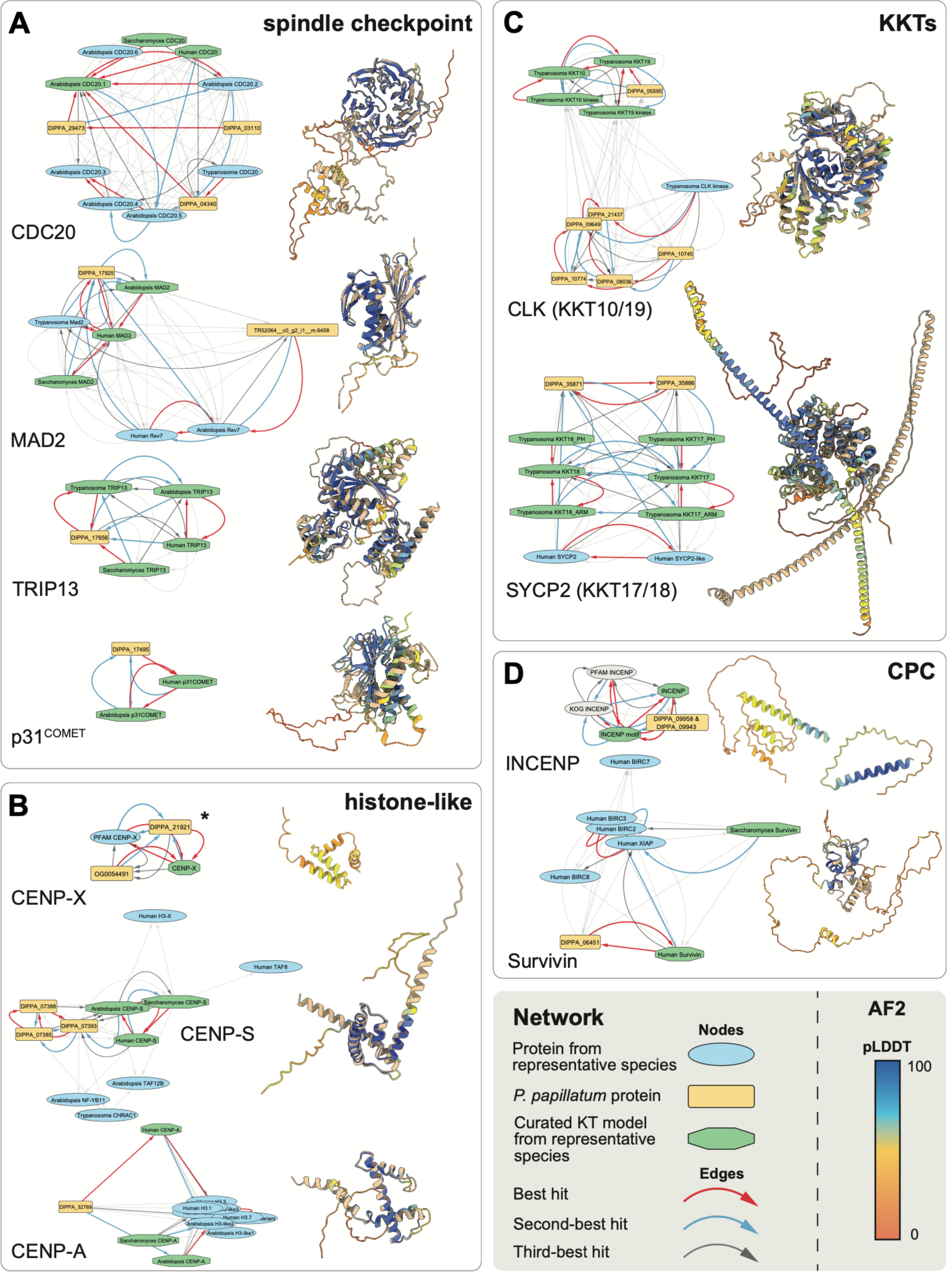
Detection of bona fide kinetochore proteins in *P. papillatum*. Network graphs showing similarity links (E-value) as inferred by Foldseek or HHsearch, displayed as edges between homologous folds depicted as nodes (left) and structural alignments as produced by Foldseek where available (right). Network graphs represent an individual Markov Clustering (MCL) cluster. Structural alignments show the predicted structure of the relevant *P. papillatum* protein placed in the cluster to the AF2 structure that was best hit in the Foldseek against the foldomes of representative species, colored in tan. Predicted protein structures are colored by pLDDT score following the standard AF2 palette. Panels represent different functional/structural aspects of the *c*KT and KKT, as indicated. *MCL clusters obtained from a network based on HHsearch homology inferences, no structural alignment is available for these models as the folds do not contain a clear structural domain. The best-predicted (highest pLDDT score) AF2 model of the *P. papillatum* accession is shown on the right.

By contrast, we were unable to identify additional *c*KT/KKT orthologs, including the principal microtubule-binding components of the *c*KT (NDC80/NUF2) and the KKT (KKT4) (**Fig. 3**), as well as other hallmark features of the inner (CENP-C) and outer *c*KT (MIS12) (**Fig. S3**). Although we found homologs for these proteins by virtue of their conserved domains e.g. the CENP-C cupin domain and the NDC80/NUF2 CH domain, these are more similar to closely-related known outgroup proteins (**Figs. 3A, S3A**). Identification of CCDC93 as an outgroup for NDC80 is in line with the LECA kinetochore reconstruction (11, 49). For other manual explorations and checks, we refer to the specific notes on each *c*KT/KKT component (**Tables S3-S5)**. Finally, we found a strong similarity link (best hit: DIPPA_32591) for KKT4, the only known microtubule-binding protein of the kinetoplastid kinetochore. However, the BRCT domain in DIPPA_32591 is a bidirectional best hit with NUP92/MLP2, a nuclear pore component in *T. brucei* (50). Indeed, our phylogenetic analysis shows that KKT4 originated through duplication of NUP92/MLP2 in the ancestor of all kinetoplastids and that DIPPA_32591 belongs to the NUP92/MLP2 family rather than KKT4 (**Figs. 3B, S4A**).

**Fig 3.**
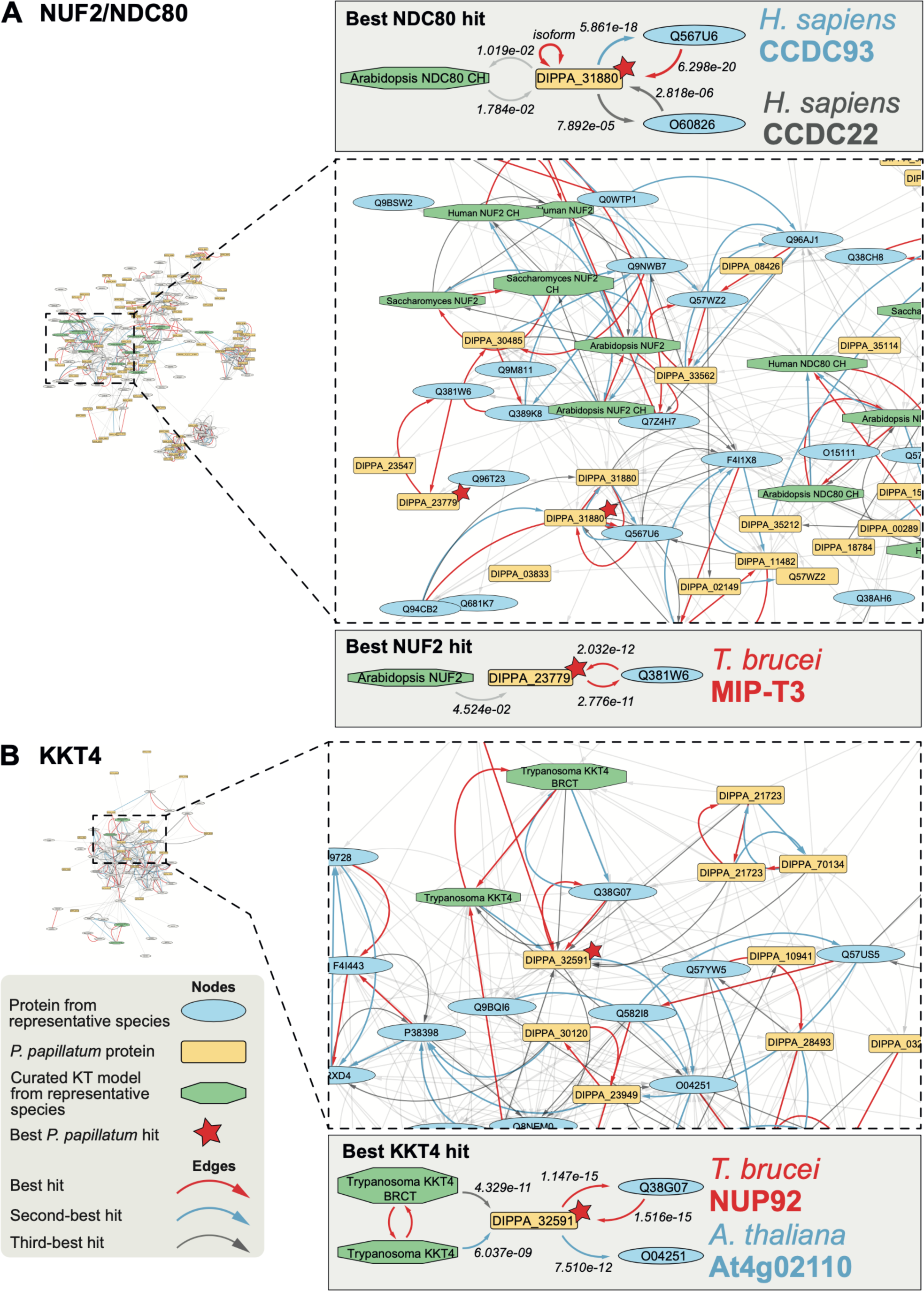
Closest homologs to NDC80/NUF2 and KKT4 are not 1-to-1 orthologs but outparalogs CCDC93/Cluap1 and NUP92, respectively. Network graphs showing homology hits as inferred by Foldseek, displayed as edges between homologous folds depicted as nodes. (**A**) All nodes connected to NUF2 & NDC80 nodes through either incoming or outgoing edges, and all nodes connected to these nodes with either directionality to the edge (i.e. all nodes until one degree of separation to a NUF2 and/or NDC80 node). Nodes are positioned according to the E-value represented by connecting edges, where shorter edges equate to lower E-value. On the left, a zoom-in of the network around the node of the *P. papillatum* accession that is the closest homolog by one of the NUF2/NDC80 query models (marked by a red star) and its closely-related hits. Schematic representations of the path connecting NDC80 (top) and NUF2 (bottom) query nodes to their respective top *P. papillatum* accession hit and its subsequent best hits are shown. (**B**) Similar to panel A, but with nodes connected to KKT4 through maximally one degree of separation.

Combined, our approaches robustly confirm the absence of a large number of components of both the *c*KT and the KKT in our diplonemid transcriptomic database and the high-quality genome of *P. papillatum* (46).

### Parallel expansions of kinetochore-related genes in Euglenozoa

In some of our clustering analyses, it proved difficult to make a confident assessment of putative orthologs solely through inspection of the network, particularly in gene families with a large amount of recent duplications. Furthermore, only the top 10 hits were considered in the network analysis, possibly missing highly-divergent orthologs. We therefore manually explored both sequence and structure-based search outputs to investigate whether our network analysis missed *c*KT/KKT orthologs.

Through careful inspection of the data, we identified a *bona fide* ortholog of the *c*KT-associated kinesin CENP-E through a bidirectional best hit (DIPPA_27648), and the closely-related KKT-associated kinesin KIN-A (DIPPA_28866 & DIPPA_16905), which appear to be part of one related family of kinesins (**Table S3C**, **Fig. S4B**). In addition, we found three BUB3/RAE1-like sequences (DIPPA_18051, DIPPA_04497 and DIPPA_14534), for which subsequent phylogenetic analysis showed that DIPPA_18051 and DIPPA_04497 are *bona fide* RAE1 orthologs, and that DIPPA_14534 branches off prior to the split of BUB3/RAE1, similar to the BUB3-like KKT15 protein (**Fig. S5A**). Based on this phylogeny, we speculate that DIPPA_14534 is most likely a BUB3 ortholog considering the presence of clear RAE1 orthologs in diplonemids and it being similarly placed outside (but closer to BUB3/RAE1) of the RAE1/BUB3 group similarly to the BUB3-like protein KKT15 (51). For the phosphatase KKIP7, we found many paralogs in Euglenozoa which arose through a complex history of duplications, with DIPPA_23494 as the 1-to-1 KKIP7 ortholog (**Fig. S5B**). Lastly, we generated phylogenetic trees for the Aurora-Polo kinases families, which were previously found to be each other’s closest paralogs in the LECA genome (52). We added the KKT2 and KKT3 kinases because these proteins also have a kinase and polo-box domain for which we found strong similarity with Polo kinases in our network analysis, similar to previous findings (23) (**Table S3**, **Fig. S6**). Diplonemid Aurora kinases appear to cluster close to those of kinetoplastids, suggesting a shared history of duplications, with a total of three Aurora kinases: Aurora-1 (Aurora B-like), Aurora-2 and Aurora-3 (**Fig. S6**). Remarkably, the kinase domains of KKT2 and KKT3 cluster at the base of the Polo kinase group closer to those of PLK4. Given that no PLK4 ortholog is present in Kinetoplastea, this might suggest that KKT2/3 are descendant from PLK4. However, the higher similarity of the polo-box of KKT2/3 with those of PLK1 and not PLK4, may well point to KKT2/3 being highly divergent Polo kinases not belonging to the PLK4 clade, and that the current position reflects extreme divergence causing long-branch attraction. We also found multiple additional parallel duplications amongst Polo kinases giving rise to distinct numbers of paralogs amongst various eukaryotic groups, including separate duplications in euglenids, diplonemids, and kinetoplastids (**Fig. S6**).

### Phylogenetic distributions of kinetochore proteins suggest distinct origins for the kinetochores in kinetoplastids, diplonemids and euglenids

We next profiled the presences and absences of *c*KT and KKT orthologs in an extended set of species (**Table S1**) belonging to the supergroup Discoba to infer the evolutionary timing of the losses, gains and transitions that occurred amongst euglenozoans, and compare these to the kinetochore compositions of *H. sapiens*, *A. thaliana,* and *S. cerevisiae* (**Fig. 4**, **Table S6**).

**Fig 4.**
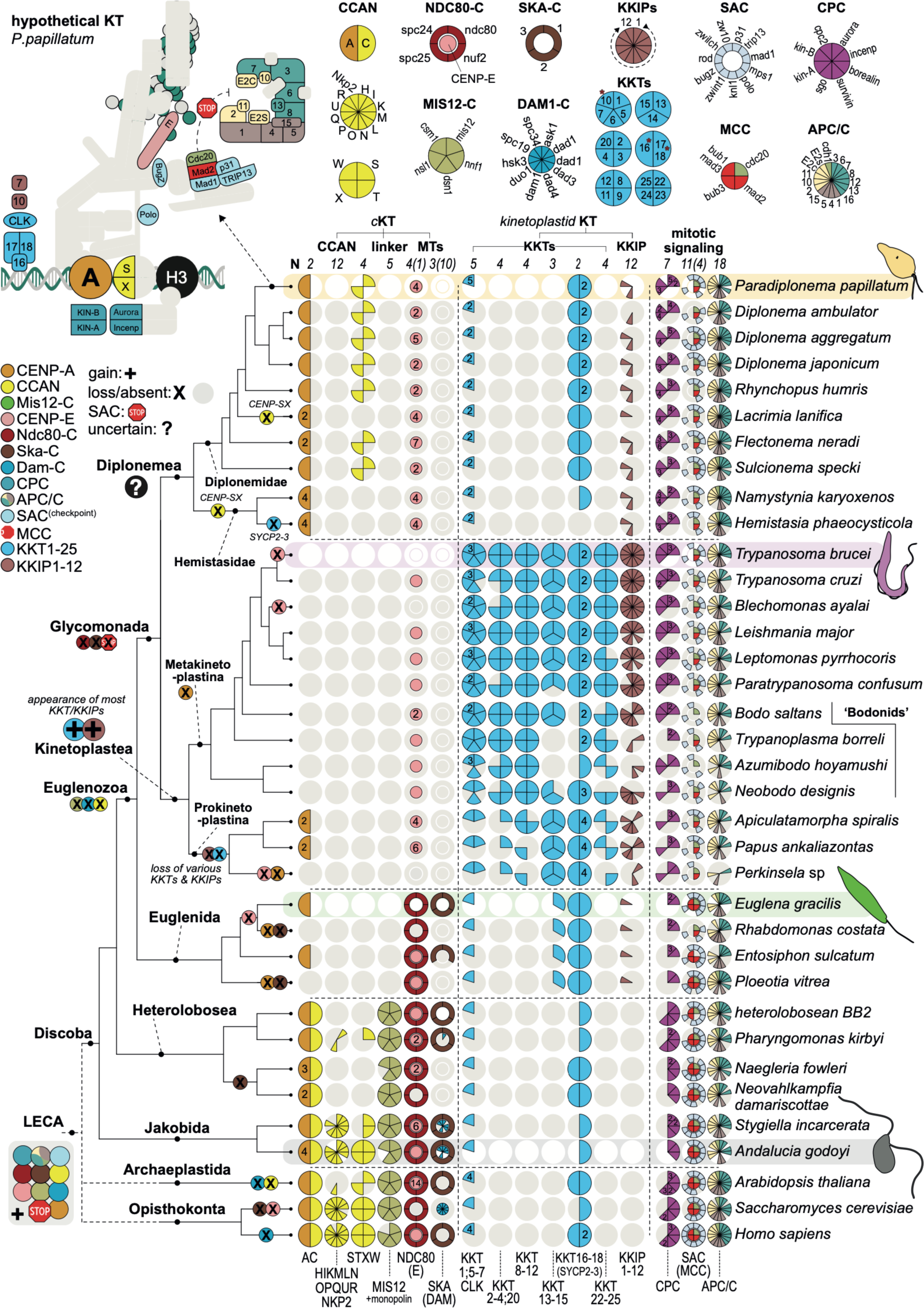
Multiple kinetochore systems in the supergroup Discoba. Presence/absence pie chart (coulson plot) matrix of both canonical as well as kinetoplastid-specific kinetochore proteins in the eukaryotic supergroup Discoba, including a projection of relevant evolutionary events for *c*KT and KKT proteins onto the tree (cartoon left). Kinetoplastea have kinetochores consisting of KKT/KKIP subunits. Euglenida have kinetochores with the conserved Ndc80 complex, CENP-A, the SKA complex and most likely an active spindle checkpoint (Bub1/BubR1, Mad2, Mad1, Bub3). Diplonemea harbor a CENP-A-like protein, only minimal numbers of KKT/KKIPs (CLK^KKT10/19^, SYCP3^KKT16^, SYCP2^KKT17/18^, KKIP7 and KKIP10) and *c*KT proteins (p31^COMET^, CENP-S/X, Mad1, Mad2). Overall, Euglenozoa appear to have lost and replaced the *c*KT in multiple steps in the different clades - see before. Ancestrally Discoba would have had a complex kinetochore system with a full CCAN (Jakobida), Mis12-C, Ndc80-C, DAM-C and SKA-C, including an active spindle checkpoint and APC/C. Top left: cartoon of conserved *c*KT/KKT-related proteins present in *P. papillatum* as projected onto the LECA *c*KT composition; grey indicates absent, coloured proteins indicate present. Top right: coulson-style plots of *c*KT/KKT complexes - below: coloured parts of the pie chart are present, light grey means these proteins are absent. Numbers indicate the number of paralogs present. Colours correspond to the cartoons of the *c*KT and KKT shown in Fig. 1. Per major taxonomic groups amongst Discoba a representative species is highlighted, including the presence of a cartoon of its cellular outline. Bottom: presence/absence of three model eukaryotes for which kinetochores have been determined: *H. sapiens, S. cerevisiae and A. thaliana*.

We found that the majority of the *c*KT is conserved in a basal group of discobans, the Jakobida (53), including the complete CCAN, MIS12, NDC80, DAM1 and SKA complexes. The presence of the CCAN in other basal lineages amongst various eukaryotic supergroups (11, 54) further bolsters the notion that the *c*KT is an ancient structure that was already present in the LECA (11). The presence of both DAM1 and SKA complexes in Jakobida, is particularly striking, as this dual presence was recently inferred to have been the case in the LECA (55). All in all, such a complex kinetochore system in a basal lineage of a supergroup highlights the notion that Discoba is one of the candidate lineages to be close(st) to the root of the eukaryotic tree of life (56). Our phylogenetic profiling shows a sequential loss of *c*KT components along the Discoba phylogeny, with first the bulk of the CCAN and the DAM1 complex being lost in Euglenozoa and Heterolobosea, followed by the gradual losses of CENP-C and MIS12 complex in Euglenozoa, and then finally the outer kinetochore (KMN & SKA complexes), components of SAC (eg. Bub(R)1) and recurrent losses of CENP-A in Diplonemea and Kinetoplastea. This pattern may be indicative of a stepwise replacement of the *c*KT by a novel kinetochore-like system in Discoba.

With respect to the KKT system, we found some new, but in general lower number of orthologs of KKT/KKIP components in the early-branching prokinetoplastids and bodonids compared to the crown trypanosomatids. Clearly, these KKT/KKIP components cannot be found outside of kinetoplastids, apart from the aforementioned CLK^KKT10/19^, SYCP2^KKT17/18^, and KKIP7 in most Discoba, as well as candidates for KKIP4, KKIP10, and KKT13 (**Fig. 4**, **Table S6**). Prokinetoplastina takes an interesting intermediate position, as they appear to have some *c*KT components, similarly to those found in diplonemids, namely p31^comet^ and CENP-A. Diplonemids additionally have a relatively complete list of the SAC and anaphase promoting complex/cyclosome (APC/C) components, with only Bub1, Cdh1, APC12, and APC16 missing. Furthermore, presence of a putative APC15 ortholog suggests that diplonemids have active SAC silencing (57). Remarkably, CENP-S/X were found in diplonemids, but not in any other euglenozoan. Such isolated presence of CENP-S/X may be explained by their role in DNA damage signalling through the Fanconi anemia pathway (58). Strikingly, however, our survey revealed the general absence of Fanconi anemia pathway components in diplonemids, suggesting that CENP-S/X might have yet another role to play.

### Endogenous tagging of putative kinetochore-related proteins in *P. papillatum*

The *in silico* analyses were followed by experiments in which we focused on five proteins. A putative CENP-A homolog (DIPPA_32769), Mad2 (DIPPA_17925), INCENP (DIPPA_09943), CLK^KKT10/19^ (DIPPA_05595), and SYCP2L1^KKT17/18^ (DIPPA_35871) were C-terminally tagged with a protein A tag (PrA) using the plasmid pDP002 that contains neomycin as a selectable marker (43). In addition, we created a cell line in which CENP-A was C-terminally tagged with V5 using a modified pDP011 plasmid (44). Following electroporation and drug selection, we assayed the expression of tagged proteins by immunoblotting using polyclonal antibodies against either protein A or the V5 tag, and typically selected three random clones for further analysis. Immunoblots of a representative clone for each tagged protein are shown (**Fig. 5**). Specific bands of the expected molecular weight were detected for each cell line, confirming the proper integration of the tags.

**Fig 5.**
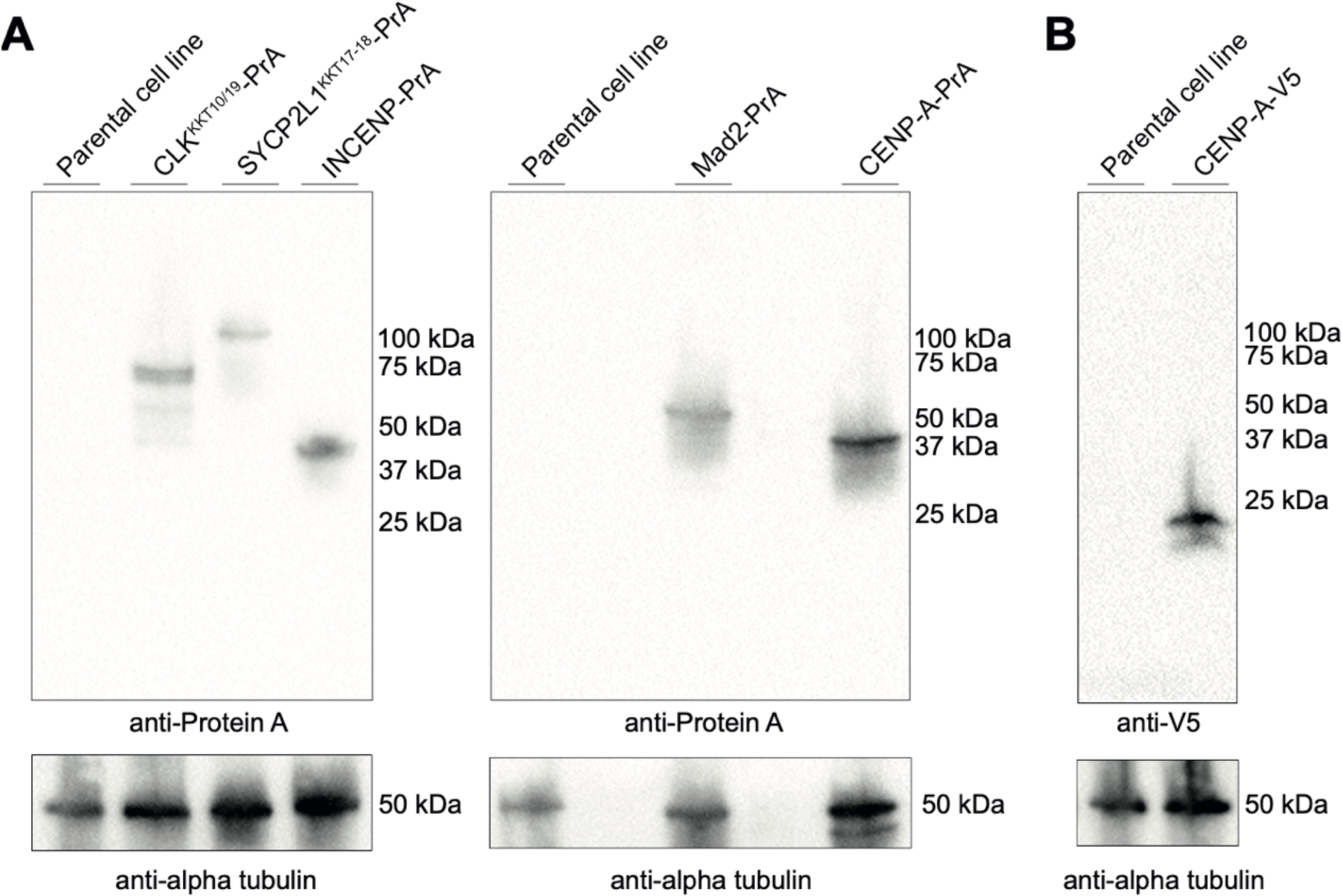
Endogenous tagging of putative mitotic proteins in *P. papillatum*. (**A**) Immunoblot analysis of *P. papillatum* cell lines expressing a Protein-A (PrA) fusion of CENP-A, INCENP, Mad2, CLK^KKT10/19^, and SYCP2L1^KKT17/18^. Anti-alpha tubulin was used as a loading control. (**B**) Western blot analysis of *P. papillatum* parental cell line and transgenic cell lines expressing CENP-A-V5.

### The CENP-A candidate forms discrete nuclear foci in *P. papillatum*

CENP-A is a centromere-specific histone H3 variant, often characterized by the longer loop 1 region that separates the helices 1 and 2 in the histone fold domain (59). Among the histone H3-like proteins in *P. papillatum*, DIPPA_32769 has many features of CENP-A, including the longer loop 1 and replacement of glutamine (Gln,Q) that is conserved in histone H3 but often changed to another amino acid in CENP-A homologs (**Fig. 6C**). In fact, our reciprocal structure-based homology search firmly placed DIPPA_32769 in a *bona fide* CENP-A cluster (**Fig. 2B**). Fluorescence and confocal microscopy analyses revealed discrete foci throughout the nucleus for both CENP-A-PrA and CENP-A-V5 (**Fig. 6A,B**). To further characterize this protein, we performed affinity purification mass spectrometry on cells expressing CENP-A-V5, using wild-type *P. papillatum* cells as a control (**Fig. S7**). Although relatively high background and lower amount of peptides precluded clear designation of strong *Pp*CENP-A interactors, we found eight homologous proteins of apparent prophage origin enriched in CENP-A-V5 pulldowns (**Table S7**), suggesting that these proteins might be associated with *Pp*CENP-A.

**Fig 6.**
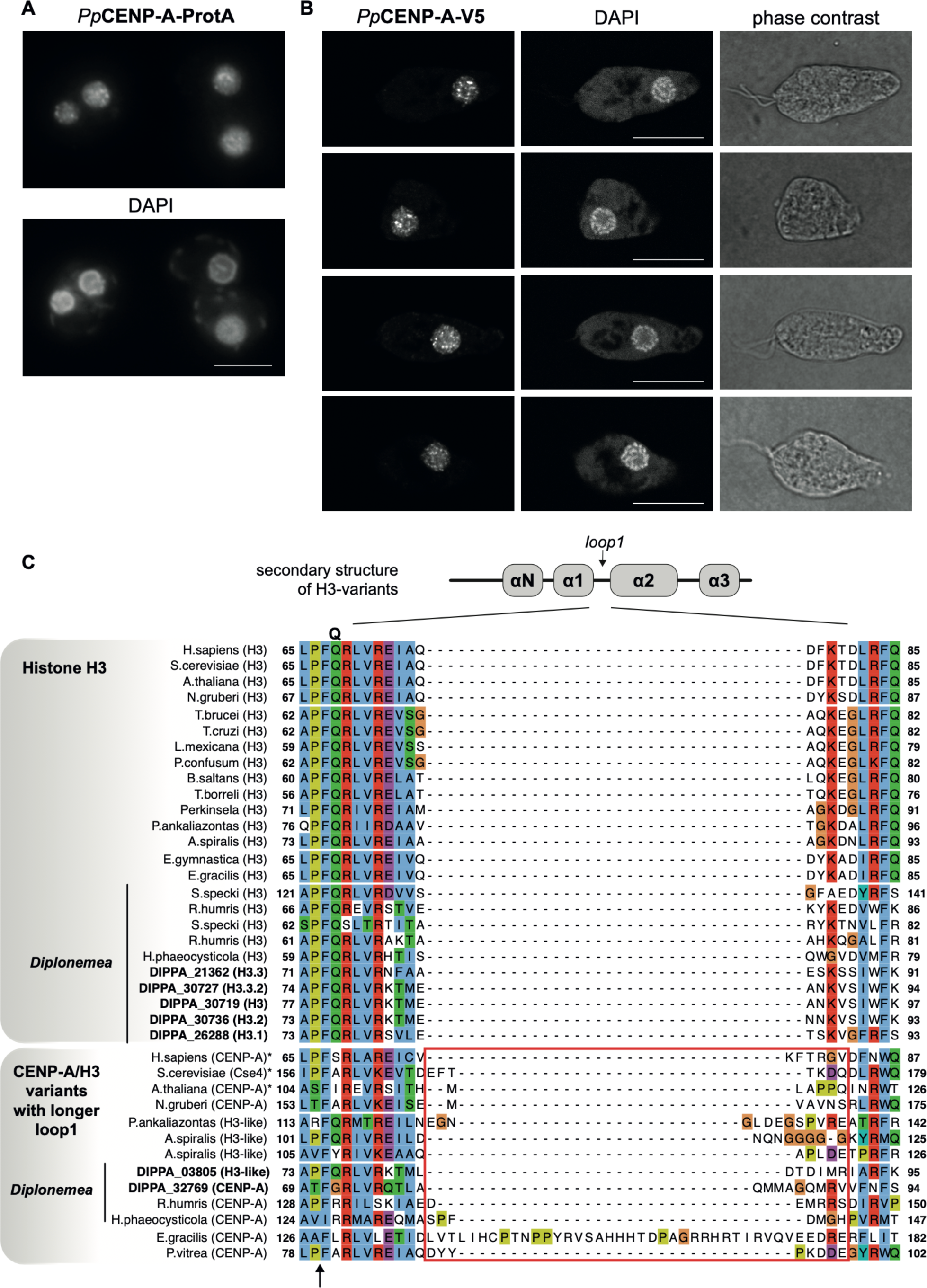
DIPPA_32769 is a putative CENP-A candidate in *P. papillatum* that forms distinct foci in the nucleus reminiscent of centromeric staining. (**A**) Immunostaining of CENP-A-PrA visualized by standard fluorescence microscopy. (**B**) Immunostaining of CENP-A-V5 visualized using a confocal microscope confirming its distinct distribution pattern in the nucleus. Single slices are shown. Image files that have full z-sections are available in Supplemental material (File S4). Scale bars: 10 μm. (**C**) Multiple sequence alignment of the CENP-A candidate in *P. papillatum* with histone H3 and CENP-A from various eukaryotes. Top: secondary structure of histone H3-like proteins. Note that DIPPA_32769 has a longer loop 1 (highlighted in the red box) as well as replacement of Gln (arrow), which are characteristic features of CENP-A. Note that DIPPA_03805 has a longer loop 1 but has Gln. CENP-A candidates are present in other diplonemids (*Rhynchopus humris* and *Hemistasia phaeocysticola*), and putatively in prokinetoplastida (*A. spiralis*).

### Diplonemids have a putative spindle assembly checkpoint system

Mad2 is a component of the spindle checkpoint conserved in many eukaryotes (12). It interacts with Cdc20, an activator of the anaphase promoting complex, which has a Mad2-binding motif (MIM) (60). Mad2 is part of the HORMA domain family, which additionally includes p31^COMET^, Hop1, and Rev7 (61). In *T. brucei* that lacks a canonical spindle checkpoint, the Mad2-like protein is associated with the basal body area and its Cdc20 lacks a Mad2-binding motif (21). Interestingly, our immunofluorescence assay revealed that Mad2 has nuclear localization in *P. papillatum* (**Fig. 7A**), implying that this protein may play a spindle checkpoint function. Consistent with this possibility, a Mad2-binding motif is present in *P. papillatum* Cdc20 (**Fig. 7A**), which is predicted by AF3 to interact with Mad2 (iPTM: 0.66). These findings suggest that a functional interaction is present between Mad2 and Cdc20 and that diplonemids likely have some form of a spindle checkpoint system.

**Fig 7.**
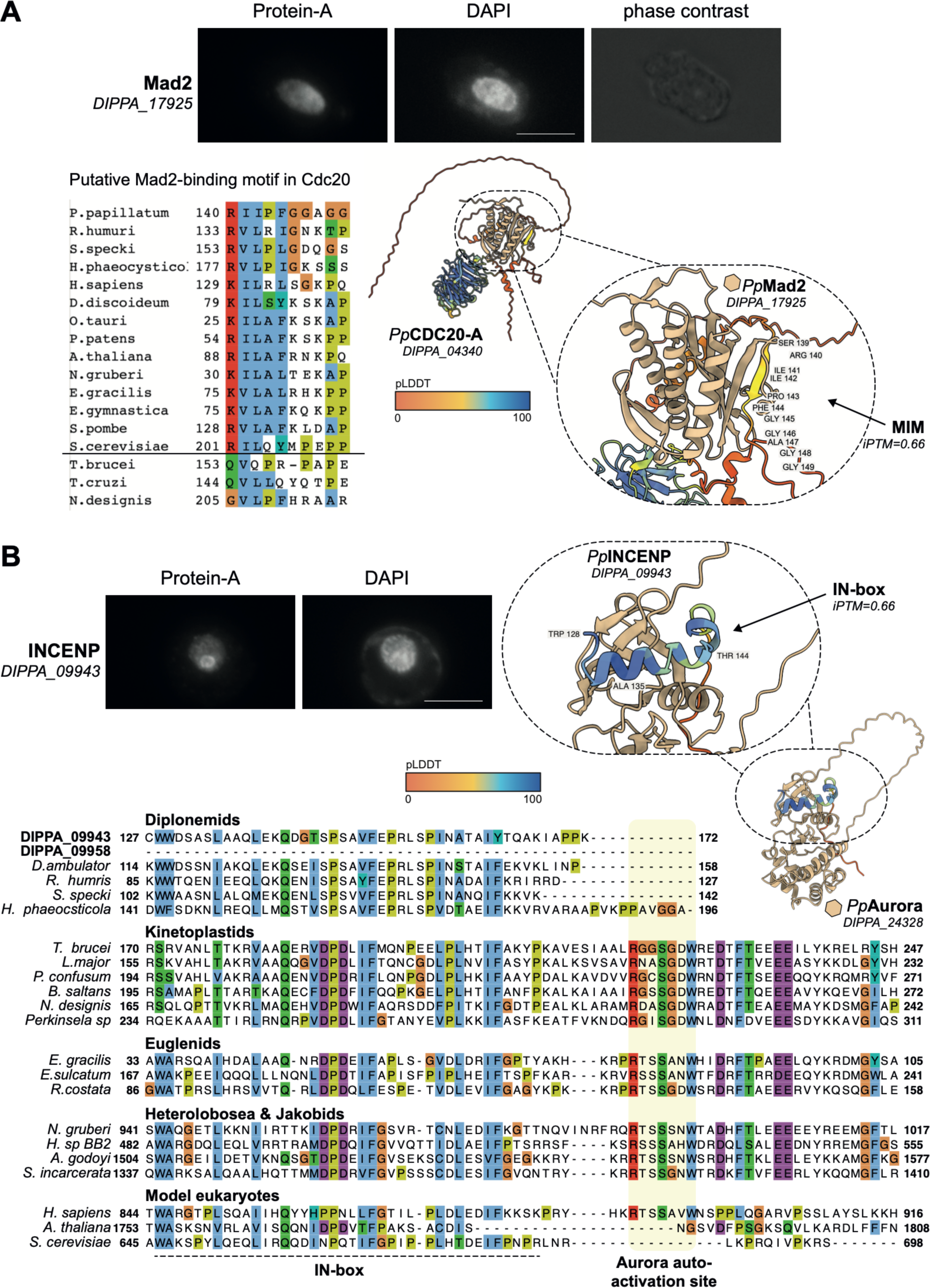
*P. papillatum* Mad2 and INCENP are nuclear proteins. **(A)** Mad2-PrA localizes in the nucleus. Scale bar: 10 μm. Cdc20 in diplonemids (*P. papillatum*, *R. humris,* and *H. phaeocysticola*), but not in kinetoplastids (*T. brucei, T. cruzi,* and *N. designis*), has a putative Mad2-binding motif. AlphaFold3 prediction of *Pp*Cdc20-A (DIPPA_04340) interaction with *Pp*Mad2 (DIPPA17925) via the predicted MIM (67), with an iPTM score of 0.66. The predicted Mad2-Cdc20 interaction indicates that a form of SAC-like signalling is active in Diplonemea. **(B)** *Pp*INCENP lacks the C-terminal cross-phosphorylation TSS site needed for proper activation for Aurora kinases in some model organisms e.g. humans. Yeasts and *Arabidopsis* appeared to have lost this motif. AF3 prediction of INCENP interaction with Aurora paralogs, pinpointed DIPPA_24328 as the most likely interaction partner (iPTM:0.66). See Fig. S9 for AF3 interaction predictions between INCENP and Aurora kinases paralogues.

### The CPC in *P. papillatum* is unconventional

The chromosomal passenger complex (CPC) is essential for orchestrating both chromosome segregation and cytokinesis (62). In most eukaryotes, the CPC consists of Aurora B kinase, INCENP, Borealin, and Survivin. However, the CPC differs from this conventional form in *T. brucei*, where no orthologs of Borealin and Survivin have been identified. Besides Aurora B-like and a highly divergent INCENP-like protein CPC1, *T. brucei* has three novel CPC components known as CPC2, KIN-A, and KIN-B (**Fig. 1C**) (63, 64). Through our bioinformatics analyses, we identified four Aurora homologs (**Fig. S6**) and putative orthologs of INCENP (DIPPA_09943 & DIPPA_09958) and Survivin (DIPPA_06451) in *P. papillatum* (**Fig. 2**). Interestingly, these CPC components have atypical features in this flagellate. The C-terminus of INCENP is defined by an IN box, which is essential for Aurora B activation (65, 66). Although orthologs of INCENP in Discoba have a highly conserved IN box, the otherwise conserved Aurora activation site located downstream of the IN box is absent in Diplonemea, with DIPPA_09958 entirely lacking the IN box (**Fig. 7B**). It is unknown to which extent this truncation in Diplonemea influences its efficacy to activate Aurora, but we note that the *S. cerevisiae* ortholog appears similarly truncated (**Fig. 7B**) (65). *P. papillatum* is the only diplonemid with two INCENP orthologs, which likely arose from a recent duplication after which DIPPA_09958 lost the IN box.

We then generated AF3 models (67) of INCENP^DIPPA_09943^ onto the four Aurora homologs in *P. papillatum* to investigate the binding of the IN box to the Aurora kinase domain and to delineate which Aurora homolog is its likely interaction partner. We find that the fold with the Aurora^DIPPA_24328^(homologous to Aurora-3/AUK3 in *T. brucei*) (**Figs. S6 and S8**) had both the highest iPTM score (0.66) as well as the highest pLDDT score in the IN box of DIPPA_09943 (**Fig. 8A**). Interestingly, INCENP^DIPPA_09943^ and Aurora^DIPPA_24328^ are predicted to interact in a similar manner as in other eukaryotes (**Fig. S8B-E**). Importantly, INCENP^DIPPA_09943^ seemingly clusters around the nucleolus (**Fig. 7B**). Given that centromeres associate near nucleoli in metazoa (68–70), the observed localization of INCENP^DIPPA_09943^ could indicate a centromere-bound state.

**Fig 8.**
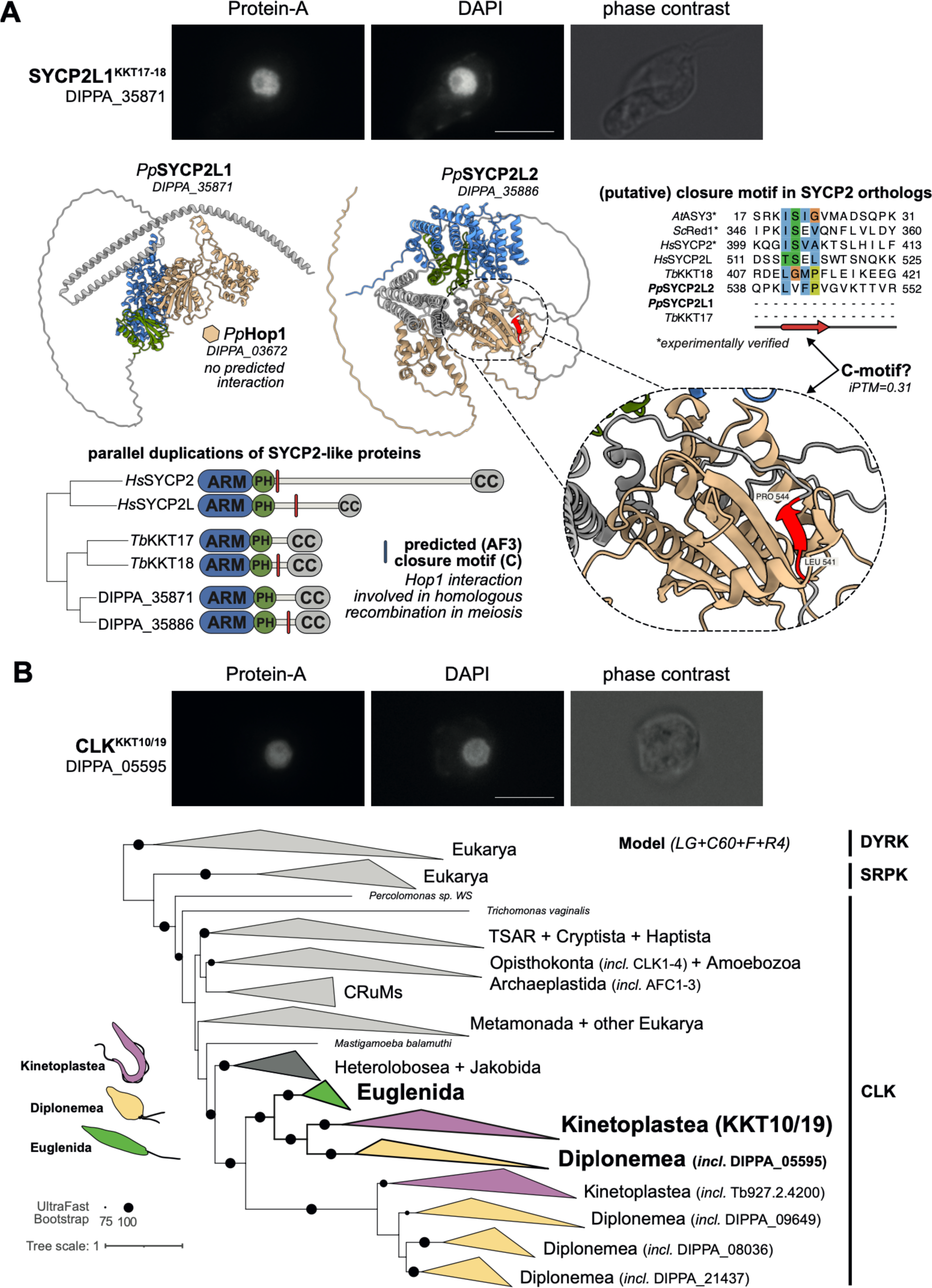
SYCP2L1^KKT17/18^ and CLK^KKT10/19^ are localized to the nucleus. (**A**) SYCP2L1^KKT17/18^ (DIPPA_35871)-PrA localizes in the nucleus. Scale bar: 10 μm. Two paralogs of SYCP2 are present in *P. papillatum*: DIPPA_35871 (SYCP2L1) and DIPPA_35886 (SYCP2L2). Right: AF3-based predictions of putative closure motifs SYCP2 orthologs that interact with the HORMA protein Hop1, showing specifically the interactions predicted for SYCP2L1 and SYCP2L2. No closure motif was found in SYCP2L1, hinting at a role for SYCP2L1 during mitosis and SYCP2L2 during meiosis (*i*PTM:0.31). Top right: alignment of verified and predicted closure motifs. **(B)** CLK^KKT10/19^ (DIPPA_05595)-PrA localizes in the nucleus. Scale bar: 10 μm. Phylogenetic tree of the CLK kinase family including both KKT10/19 and the closest paralog in *P. papillatum* (DIPPA_05595).

Unexpectedly, we identified DIPPA_06451 as a BIR domain-containing protein that has a bidirectional best hit in our Foldseek analysis to the BIR domain found exclusively in the opisthokont Survivin orthologs. However, it has been shown recently that the BIR domain is not an ancestral feature of Survivin, as it is only found in Opisthokonta (71). Furthermore, we do not find homology of the N-terminus of DIPPA_06451 to the N-terminal coil that is the defining feature of Survivin orthologs across eukaryotes (71). DIPPA_06451 is therefore unlikely a true Survivin ortholog. This reiterates the importance of looking beyond a bidirectional best hit, which is often considered sufficient to classify orthologs.

Finally, we identified three putative KIN-A orthologs (DIPPA_17532, DIPPA_28866, DIPPA_16905; **Fig. S5B**) through a phylogenetic analysis of kinesin-related proteins. Our phylogeny places KIN-A and KIN-B as early-branching clades nested within a larger clade of CENP-E orthologs. However, the KIN-B clade has a long branch, which is relatively poorly-supported by bootstrap scores (54/100). By contrast, KIN-A is placed within the CENP-E group with a high support from bootstraps (96/100), providing a robust phylogenetic support for the notion that KIN-A might be a highly-divergent CENP-E ortholog. It is important to note that this tree topology is unlikely due to under-sampling of KIN-A-related kinesins, as we have specifically included additional sequences from dedicated KIN-A and KIN-B HMM searches. Furthermore, a profile-versus-profile search shows that CENP-E is the most significant hit for KIN-A, and KIN-A is among the top hits when the CENP-E profile is taken as a query (**Table S4**).

### CLK^KKT10/19^ and SYCP2L1^KKT17/18^ localize in the nucleus

We next examined the localization of proteins that have similarity to kinetoplastid kinetochore proteins, but most likely perform other functions in *P. papillatum*, namely CLK^KKT10/19^ and SYCP2L1^KKT17/18^ (**Fig. 8**). Unlike CENP-A, we did not observe discrete dot signals for CLK^KKT10/19^ or SYCP2L1^KKT17/18^, although both are localized in the nucleus. It therefore remains unclear whether they are kinetochore proteins in *P. papillatum*. It is noteworthy that expression of the SYCP2L1 protein was observed in our mitotic culture despite the fact that SYCP2 is typically a meiosis-specific protein in other eukaryotes (72). Besides SYCP2L1, diplonemids have another paralog (SYCP2L2; DIPPA_35886), similarly to KKT17/18 in kinetoplastids and SYCP2/SYCP2L in vertebrates (31). SYCP2-like proteins in part function during meiosis by specifically recruiting Hop1/HORMAD to the synaptonemal complex. Indeed, we find a predicted interaction motif (iPTM: 0.31), termed closure motif, in SYCP2L2 of *P. papillatum*, suggesting that this paralog might perform a function during meiosis (**Figs. 8A** and **S9**). DIPPA_25410 is a candidate SYCP3-like protein that may interact with the SYCP2L1-2 paralogs. Like SYCP2L1, expression of SYCP3 (DIPPA_25410) is found in the background of the CENP-A-V5 pulldown (**Table S7**).

The CLK kinase family of splicing proteins operate at the kinetoplastid kinetochore as KKT10/19 (73). Since we observed that this gene family underwent specific expansions in diplonemids, akin to humans and plants (74, 75), we wondered whether KKT10/19 were phylogenetically more associated to one particular CLK in these protists. Previous studies identified DIPPA_05595 as a KKT10/19 ortholog (29). Hence, we performed more extensive analyses that included additional eukaryotic lineages as well as available discoban lineages (**Fig. 8B**). We found that KKT10/19 and DIPPA_05595 follow the species tree and are part of a seemingly slower evolving clade of the CLKs in Euglenozoa, with a split at the base of Euglenozoa subsequently giving rise to three and one diplonemid and kinetoplastid CLK, respectively (**Fig. 8B**).

Taken together, our deep homology searches did not reveal orthologs for most of the kinetoplastid or canonical kinetochore subunits in Diplonemea, suggesting that a functionally analogous structure must be responsible for facilitating chromosome segregation. This as-yet elusive kinetochore structure possibly consists of a novel type of kinetochore proteins.

## DISCUSSION

Presently, the only group of euglenozoans where the kinetochore has been experimentally studied in detail are the trypanosomatid flagellates, which include human pathogens such as *T. brucei*, *T. cruzi*, and *Leishmania* spp. (22, 24, 76, 77). Unique KKT proteins identified in *T. brucei* are conserved even in early-diverging prokinetoplastids (29), meaning that the KKT-based system likely already emerged in the last common ancestor of Kinetoplastea. To gain insights into the origin of the unique kinetoplastid kinetochore, and to uncover a putative novel kinetochore system, we sought to examine and identify kinetochore proteins in diplonemids, the sister clade of kinetoplastids (29, 33, 34). To this end, we developed a highly sensitive pipeline tailored to identify even highly-divergent orthologs of the *c*KT and KKT, by leveraging sequence- and structure-based deep homology detection methods, using for instance a near-complete predicted foldome of the model diplonemid *P. papillatum*, and combining these with extensive phylogenetic analysis. Our approach has led to the identification of several highly-divergent *c*KT orthologs, including a p31^COMET^, Mad1 and Bub3 ortholog in *P. papillatum*, establishing the validity of our approaches. Yet, even these state-of-the-art methods were unable to identify additional *c*KT/KKT orthologs in *P. papillatum* and other diplonemids. In particular, they apparently do not have any of the structural kinetochore components of these systems, making it highly unlikely that these closest relatives of kinetoplastids possess any type of the known kinetochore repertoire.

Sensitivity of bioinformatics approaches to detect highly divergent homologs remains a recurring theme. Our previous efforts to characterize the evolution of kinetochore compositions across eukaryotes using iterative HMM searches suggested extensive or complete loss of core subunits not only in the euglenozoan lineages, but also in Metamonada and Alveolata (10). Although for some species like the metamonad *Carpediemonas membranifera* the widespread loss of ancestral kinetochore subunits appears to hold (19), recent work in the apicomplexan parasite *Plasmodium berghei* revealed that a combination of advanced biochemistry (BioID), profile-vs-profile HMM searches and AF2-based structure prediction was paramount to uncover extremely divergent subunits, that in hindsight appeared to be conventional counterparts (18). Our approach adds a full structural search of the predicted foldome of the model diplonemid *P. papillatum* using AlphaFold2 and Foldseek and includes high-resolution orthologous group definitions for 11 diplonemid species to strengthen profile-vs-profile HMM searches. In addition, we used standard HMM searches without heuristic filters, which we recently found out to yield more *bona fide* orthologs, though at the cost of higher computational time. To our knowledge, we here performed the most extensive set of computational analyses to disprove the presence of any protein with similarity to both canonical and kinetoplastid kinetochore subunits, or any eukaryotic protein for that matter. A priori we can never fully exclude missed detection of orthologs. As an example, our recent work revealed the likely orthology of KKT14-15 with Bub1-Bub3, which was previously missed using standard blast and HMM searches (51). In light of the ever increasing sensitivity of tools to detect highly divergent homologs, scepticism towards bold claims of protein/gene absence appear justified – and perhaps the null hypothesis for any absence should be a detection problem. We therefore strongly advocate to any who put forward claims of surprising losses, to follow our approach and to go to great lengths to disprove the presence of highly divergent orthologs.

The only canonical structural kinetochore candidate identifiable in diplonemids is CENP-A, a variant histone that replaces the canonical histone H3 at centromeres. Based on the findings that the histone H3 variant in *P. papillatum* has many features of CENP-A and localizes to discrete foci in the nucleus, we deem it most likely that it is a true CENP-A homolog. Early studies using electron microscopy found numerous short condensed chromosomes in the interphase nucleus of *P. papillatum* and *Diplonema ambulator* (78, 79). More recently, using expansion microscopy of *P. papillatum* (46) and transmission electron microscopy of *Lacrimia vacuolata* (39), we were able to visualize sub-nuclear structures, namely the large centrally located nucleolus, numerous peripheral concentric-oriented and condensed chromosomes and one or two small electron-dense spherical bodies. Consistent with a high number of chromosomes in diplonemid nuclei, confocal microscopy allowed us to discern more than 20 CENP-A-V5 foci. However, it is important to note that we do not have a kinetochore marker to prove that these signals represent true kinetochore localization and, notably, CENP-A is not the only divergent H3 variant that localizes into the nuclear foci. For example, there is an H3 variant in *T. brucei* which has been shown to localize to telomeric foci (80). Nonetheless, formation of discrete nuclear foci, together with its previously noted structural and sequence features (**Fig. 6**), suggest that DIPPA_32769 is a true CENP-A ortholog in *P. papillatum*, indicating presence of the *c*KT nucleosomal cornerstone.

We identified numerous key components of a putative spindle assembly checkpoint in *P. papillatum*, such as Mad2, Cdc20 and a Bub3 candidate. However, members of the central checkpoint protein family Bub(R)1/Mad3 were not detected. It is intriguing that *P. papillatum* has a Mad2-like protein that localizes in the nucleus, unlike *T. brucei* Mad2 that localizes near the basal body (21). Together with the finding that *P. papillatum* Cdc20 has a conserved Mad2-binding motif, it is likely that Mad2 interacts with Cdc20 to regulate APC/C activity. Interestingly, diplonemids and the early-branching prokinetoplastids encode p31^COMET^, which in other eukaryotes is involved in the regulation of the open and closed forms of Mad2 and other HORMA domain proteins (81). While it was thought that Mad1, a kinetochore recruitment factor for Mad2, would be absent from kinetoplastids, our iterative HMM searches in combination with Foldseek of AF2-predicted folds revealed the presence of an RWD domain reminiscent of Mad1 in the *T. brucei* protein LAP71, which was recently described to be a nuclear pore protein (82). Candidate orthologs in diplonemids were also found, including the *P. papillatum* protein DIPPA_30775, although a putative MIM could not be discerned. It is possible that these Mad1-like proteins have other functions at the nuclear pore complex in kinetoplastids and diplonemids. In any case, it will be important to examine whether *P. papillatum* has a functional spindle assembly checkpoint and delays cell cycle progression in response to treatment with microtubule poisons.

In addition to the absence of the bulk of the *c*KT, we also did not find critical components of the KKT. Instead, we only found putative candidates for proteins that are broadly conserved amongst eukaryotes with non-kinetochore functions (e.g. CLK and SYCP2) (29, 31, 83). Hence, these proteins most likely also have non-kinetochore functions in kinetoplastids and diplonemids. In traditional model eukaryotes such as humans and *Drosophila melanogaster*, CLK kinases do not localize at kinetochores but are instead involved in the regulation of splicing (84). Similarly, KKT16/17/18 have a combination of Armadillo, PH and coiled-coil domains, which are found in the broadly conserved synaptonemal complex proteins SYCP2 and SYCP3 (31). The facts that euglenids have homologs of the canonical kinetochore proteins and that additional orthologs of the kinetoplastid-specific KKT proteins have not been identified suggest that euglenids most likely have canonical kinetochores and that their CLK^KKT10/19^ and SYCP2/3^KKT16/17/18^ proteins have non-kinetochore functions. Although these proteins localize in the nucleus of *P. papillatum*, further studies are needed to reveal their functions.

The cell lines prepared in this study will serve as an invaluable resource in further dissection of the seemingly unique *P. papillatum* kinetochore whose composition remains unknown. Although the resolution of the performed pull-down experiments with CENP-A-V5 was generally low, our experiments hint at a putative interaction with eight proteins that all bear a homologous coiled-coil domain that is of prophage origin. These proteins should be characterized in the future, as they might play a role in the diplonemid kinetochore.

Diplonemids are among the most abundant and diverse species in the oceans (35–38), yet we know very little about their basic cell biology (44–47). Here we have shown that these heterotrophic flagellates must segregate their chromosomes in a process involving an as-yet elusive kinetochore-like structure, a core apparatus in the eukaryotic cell cycle. Moreover, as a sister clade to the (mostly) parasitic kinetoplastids, which are endowed with a number of unique features (85), diplonemid cell biology may shed light on their evolution as well as increase our understanding of one of the most abundant yet deeply understudied group of marine eukaryotes.

## MATERIALS AND METHODS

### Diplonemid Transcriptomic Dataset

For this study a set of predicted proteomes of 11 Diplonemid species were used; 1 genome (*P. papillatum*) (46) and 10 transcriptomes (29, 46, 86–88) (**Table S1**). For de novo assembly of RNA-Seq datasets, Trinity v2.2.0 software was used with default parameters (https://github.com/trinityrnaseq/trinityrnaseq) (89). TransDecoder v5.5.0 (Haas B.J., https://github.com/TransDecoder/TransDecoder) was used to generate proteomes using ‘Universal’ genetic code. Proteins were then clustered with CD-HIT v4.8.1 at 99% identity (https://github.com/weizhongli/cdhit/) (90) to reduce redundancy and proteome complexity. *Paradiplonema papillatum* identifiers were translated to the following: ‘Paradiplonema_papillatum_xxxxxx, where x is a six digit number (**Table S2**).

### AlphaFold2 Foldome for *P. papillatum*

The structures of all predicted protein sequences from the recently-sequenced *P. papillatum* genome were predicted with AlphaFold2 (AF2) (46, 91). It has been shown that supplying multiple sequence alignments (MSAs) from closely-related species can improve the confidence and likely accuracy of AF2 models, especially for sequences from divergent and underrepresented taxa (92). MSAs were generated using the MMseqs package (93), where each *P. papillatum* sequence was clustered and aligned to sequences in a database comprised of the EukProt v3 dataset (94), the discoba-specific dataset by (92) and our own set of predicted diplonemid proteomes (see above). The AF2 algorithm was run on these MSAs using the ColabFold implementation (95). An upper limit of 3,000 amino acids per *P. papillatum* amino acid sequence was imposed for computational tractability, resulting in a total of 41,990 predicted structures. The AF2 predictions with the highest average pLDDT per protein (rank 1), and all other related files to the AF2 predictions can be found in **File S3**.

### Deep homology detection protocols

A three-pronged highly-sensitive homology detection was used consisting of (i) a unidirectional profile Hidden Markov Model (HMM) search of known kinetochore components against the *P. papillatum* proteome, (ii) an extensive profile-vs-profile HMM search, and (iii) an extensive three-dimensional protein structure-based search. A graphic overview of the pipeline for the Diplonemea dataset generation and the HHsearch/AF2 strategy can be found in **Figs. S1** and **S2**. Raw files pertaining to these searches can found in **File S3**.

#### Unidirectional Hidden Markov Model searches without heuristic filters

Curated multiple sequence alignments of both full-length and/or domains and motifs for 206 canonical kinetochore (KT) orthologs/outparalogs (11), including highly divergent apicomplexan kinetochores (18), as well as 36 kinetoplastid-specific kinetochore (interacting) (KKT/KKIP) proteins, were converted into profile HMMs using the *hmmbuild* from HMMer package (v3.3, custom settings) (96). To distinguish from previous unidirectional profile HMM approaches (29), searches were performed with *hmmsearch* without any of the standard heuristic filters (option –max) that are commonly in place to maintain reasonable computational tractability in larger sequence databases. Surprisingly, we have found that turning these heuristic filters off regularly reveals high scoring ‘hits’ of highly-divergent homologs. ***Profile-versus-Profile -HMM*** OrthoFinder (v2.5.5, -S blast; otherwise default settings) was used to generate orthologous groups (OGs) from the predicted proteomes in our diplonemid set (see above) (97). OGs were aligned with MAFFT (v7.520, E-INS-i) (98). Using the HHsuite package (v3, standard settings), a consensus sequence was derived for each alignment (.a3m format) in order to generate secondary structure decorated profile HMMs (.hhm format), which were collated in a custom HHsearch database (99). The same procedure was followed for our manually curated set of known kinetochore proteins (see above) and queried using the hhsearch tool against the Diplonemid database. First, our manually-curated set of KT/KKT/KKIP/AKiT alignments was queried against the Diplonemid OG database. The top 10 hits for each of these initial searches were taken and were subsequently queried against the KT/KKT database, the Diplonemid database, as well as against the COG/KOG and PFAM precomputed hh-suite database (https://www.user.gwdguser.de/~compbiol/data/hhsuite/databases/hhsuite_dbs/, last accessed 01-04-2024) to provide a comprehensive and high-quality background set of profiles. This search step is considered the ‘forward search’. In turn, the top 10 hits for each of these forward searches was again queried against the same set of databases in a step considered the ‘reverse search’. Finally, the top 10 hits from all forward and reverse searches were parsed and compiled into a dataset that describes the relations of the queries and top 10 targets of each performed search. This dataset was analyzed using Cystoscape software v.3.10.2 (100). Markov Clustering (MCL) was performed on the dataset to identify clusters of nodes, which typically are most strongly interconnected amongst each other and show similar levels of connectivity to other regions in the network, which is indicative of a closely-related phylogenetic affiliation relative to other nodes in the network. MCL was performed with -log(E-value) taken as input array for edge weights and an inflation parameter of *I* = 2. The MCL network can be found in **File S1**.

#### 3D protein structure comparison

Manually-curated AF2 models (downloaded from uniprot, last accessed 25-03-2024) of known KT/KKT/KKIP proteins from four representative species: *Arabidopsis thaliana, Homo sapiens, Saccharomyces cerevisiae,* and *Trypanosoma brucei* (91) were searched against our proteome-wide predicted structures for *P. papillatum* using Foldseek (101). Both full-length and domain-specific structures were queried. First, an initial search was performed against only the *P. papillatum* AF2 foldome, from which the top 10 hits were taken as query for a ‘forward’ search against the complete database, comprising of the curated *c*KT/KKT/KKIP folds, the *P. papillatum* AF2 foldome and the complete foldomes of the representative species. Finally, the top 10 hits for each ‘forward’ query were taken and were queried again against the complete database in a ‘reverse’ search step. The top 10 hits for all forward and reverse searches were compiled into a dataset which was analyzed with the Cytoscape software, primarily through the MCL algorithm implementation. MCL was performed with -log(E-value) taken as input array for edge weights and an inflation parameter of *I* = 7. The MCL network can be found in **File S2**.

### AlphaFold3 protein-protein interaction modelling

To structurally model protein-protein interactions, AlphaFold3 was used through its online implementation at alphafoldserver.com (67), last accessed 13-7-2024.

### Phylogenetic analyses

To delineate orthologs from homologs in select cases, phylogenetic analyses were performed for several orthologous groups as follows. First, similarity searches were performed using a combination of hmmsearch and jackhmmer from the HMMER package (v.3.1b2). These searches were conducted against our in-house database (**Table S1**) and a eukaryote-wide selection of proteomes as used by De Potter et al (102). The obtained homologs were then aligned with MAFFT E-INS-i or L-INS-i (98). Trimming of the multiple sequence alignment (MSA) was performed using trimAl. Sites were removed with the criterion <= 10% or <= 30% occupancy, and finally, poorly-aligned sequences were manually discarded. A phylogenetic tree was then inferred using IQ-TREE (103) with the best-fitting substitution model as chosen by ModelFinder (104), which in all cases converged on LG+C60+F+R4. In all cases the top 500 from our searches were included in the phylogenetic analyses. For BUB3, we specifically added a known outgroup for RAE1/BUB3: WDR74 (51). A set of WDR74 ortholog sequences was downloaded from the eggNOG database (KOG3881; v5). These WDR74 sequences were combined with manually-curated MSAs of BUB3, RAE1 and KKT15. Then, the Diplonemea sequences that were assigned to the same OrthoFinder orthologous group as our candidate BUB3 sequence were also added to this set of sequences. For Aurora/Polo/KKT2/3, we included previously established Aurora and Polo clusters (11), including the LECA kinase OG PLK, Aurora and PLK4, PLK and PLK4 sharing the most recent duplication, and Aurora being an outgroup.

### Endogenous C-terminal tagging of CENP-A, Mad2, INCENP, CLK^KKT10/19^, and SYCP2L1^KKT17/18^

Parts of the open reading frame (ORF) and 3’ untranslated region (UTR) of *P. papillatum* genomic DNA were amplified using specific primers containing sequences overlapping with the protein A-neomycin cassette of the plasmid pDP002 in order to endogenously tag CENP-A, CLK^KKT10/19^, SYCP2L1^KKT17/18^, INCENP and Mad2 with a C-terminal protein A-tag. The pDP002 plasmid was used as a template for amplification of the protein A tag and the downstream Neo^R^ marker. To ligate all three fragments together, Phusion polymerase (NEB) was applied in a nested PCR using these fragments as a template. Finally, 3–5 μg of the nested PCR product was electroporated into *P. papillatum* as described elsewhere (43). Similarly, the 3xV5 and Hygromycin^R^ cassette was amplified from a slightly modified pDP011 plasmid (44), named pDP011-A (for primers see **Table S8**). The PCR product was A-tailed and cloned into pCR 2.1-TOPO (ThermoFisher) or alternatively gel-purified, ethanol-precipitated and used directly for transfections. Restriction enzyme digestion and sequencing were used to confirm the correct structure of the generated plasmid (if applicable). To release the tagging segment, 10 µg of the final plasmid was cut with EcoRI (NEB), ethanol precipitated, resuspended in 10 µl of water, and used to transfect *P. papillatum* as described elsewhere (43).

### Electroporation of *P. papillatum*

Amaxa Nucleofector II was used to transform a total of 5 x 10^7^ cells, as previously described (41, 42). Clones were selected in 24-well plates at 27 °C using varied doses of G418 for protein-A tagging (25–80 µg/mL) and of hygromycin for V5-tagging (100–225 µg/mL). Successful transfectants appeared after 2 weeks of selection. Before being tested by western blot, each clone was expanded to a volume of 10 mL and cultured for up to 2 to 3 weeks.

### Immunofluorescence

A 5 to 10 mL log phase culture was centrifuged at 1000 x g for 5 min. Cells were fixed in 4% paraformaldehyde in seawater at room temperature for 30 min. The fixative was washed out of the cells with seawater and 1 x PBS (<u>P</u>hosphate <u>b</u>uffered <u>s</u>aline) (1:1). The cells were rinsed once more in 1 x PBS before being spotted on a gelatin-coated slide, and permeabilized for 20 min for antibody labelling in 100% ice-cold methanol. Throughout the procedure, the slides were stored in a humidified chamber, washed in PBS after 20 min. and blocked with 5% milk in PBS-T (PBS with 0.05% Tween) for 45 min. After removing the blocking solution, the cells were again washed in 1 x PBS. The primary antibody (anti-Protein A; Sigma, 1:2,000) was applied on slides covered with parafilm and incubated overnight at 4°C. After removing the primary antibody, the slides were washed three times with 1 x PBS. The secondary antibody (AlexaFluor555-goat-anti-rabbit; Invitrogen, 1:1,000) was added and the slides were incubated for 1 hour in the dark at room temperature, covered with parafilm. Finally, the slides were washed in 1 x PBS and covered with 4’,6-diamidino-2-phenylindole (DAPI) containing the antifade reagent ProlongGold (Life Technologies). Images for Protein A fusion proteins were obtained using camera Olympus DP73 (Axioplan 2 imaging). CENP-A-V5 images were acquired on a FV3000 Olympus confocal microscope with an HCPL apochromatic 100x/NA 1.40 oil immersion objective. Excitation was performed with a 405 nm diode laser (50 mW) in the case of DAPI and in the case of Red channel which is Alexa Fluor 555 (543 nm, 555-615 nm); emissions were detected using hybrid detectors (HyD). Z-stacks were acquired with a step size of 100 nm (without averaging). The pixel size and dwell time were between 52 and 97 nm and 400 ns, respectively. The size of the pinhole was adjusted based on the signal strength but was typically around 0.4 Airy unit to improve resolution.

### Western blotting

Protein samples from 5 x 10^5^ cells were prepared by pelleting, resuspended in 25 µl of 2 x SDS sample buffer, separated on 4-20 % Mini-protein TGX stain-free gels (Bio-Rad), and subsequently transferred to a PVDF membrane. After blocking with 5% milk in PBS-T for at least 30 min at room temperature, the membrane was incubated with an anti-protein A (Sigma, P3775; used at 1:10,000 dilution) or anti-V5 antibody (Merck, V8137; used at 1:1,000 dilution) in 5% milk in PBS-T overnight at 4 °C. After 3 washes in PBS-T, the membrane was incubated with anti-rabbit-HRP (Merck; 1:1,000) and incubated at room temperature for 1 hour. The membrane was then washed 3x in PBS-T and the signal was developed using Clarity Western ECL Substrate (Bio-Rad). The mouse anti-alpha tubulin antibody (Merck, T9026; 1:10,000) was used as a loading control.

### Immunoprecipitation

Approximately 5 x 10^8^ cells expressing V5-tagged CENP-A proteins, as well as the wild-type control cells were grown in Diplonema growth media with the appropriate selection antibiotic. Cells were harvested at 1,000 g for 10 min, lysed using ice cold IPP150 buffer (10 mM Tris-HCl, pH 6.8, 150 mM NaCl, 0.1% Igepal, CA-630) supplemented with 1x complete EDTA-free protease inhibitors (Sigma-Aldrich, 11873580001) and 5x passed through a 30 gauge needle. The cell lysate was cleared twice by centrifugation (12,000 g, 10 min, 4 °C). Fifty µl of V5-Trap magnetic particles (M-270) (Chromotek, v5td-100) was added to the cleared cell lysate and rotated at 4°C for 2 to 3 hours. The beads were washed 3x with the IPP150 buffer containing detergent and then twice without the detergent. Bound proteins with beads were processed for immunoblotting and mass spectrometry.

### Mass spectrometry analysis

For mass spectrometry sample prep, on-bead digestion protocol was used. Briefly, washed beads were resuspended in sodium deoxycholate (SDC, final concentration 1% [w/v] in 100 mM TEAB, triethylammonium bicarbonate), reduced with 5 mM TCEP [tris(2-carboxyethyl)phosphine], alkylated with 10 mM MMTS (S-methyl methanethiosulfonate) and digested sequentially with Lys-C and trypsin. SDC was removed by extraction with ethylacetate saturated with water (105). Samples were desalted on Empore C18 columns, dried in a speedvac, and dissolved in 0.1% TFA + 2% acetonitrile. Desalted peptide digests were separated on a 50 cm C18 column using 60 min elution gradient and analyzed in a DDA mode on an Orbitrap Exploris 480 (Thermo Fisher Scientific) mass spectrometer equipped with the FAIMS unit operated at -40 and -60 V CVs.

### Data processing

The resulting raw files were converted to mzXML files each containing separate compensation voltage (CV, -40, -60 V) scans using FAIMS MzXML Generator (release 1.1.8003, https://github.com/PNNL-Comp-Mass-Spec/FAIMS-MzXML-Generator). MzXML files were analyzed in MaxQuant (v. 2.2.0.0) (106) with label-free quantification (LFQ) algorithm MaxLFQ and match between runs feature activated. FDR was set as 0.01 at all levels. A custom protein database of *P. papillatum* proteins (43,871 sequences) supplemented with frequently observed contaminants was used for protein identification. MMTS alkylated cysteine was selected as a fixed modification (Methylthio (C), composition: H(2) C S, +45.988). Variable modifications were Oxidation (M) and Acetyl (Protein N-term). Downstream processing of the proteinGroups.txt file was performed in Perseus (v2.0.6.0) (107).

## ACKNOWLEDGEMENTS

We acknowledge support from the Czech Grant Agency 23-06479X (to J.L.), and 23-07674S (to C.B.), a Veni Fellowship from the Dutch Research Council (NWO, VI.Veni.202.223) (to E.C.T.), a Wellcome Discovery Award (227243/Z/23/Z), and a Centre Core Grant to the Wellcome Trust Centre for Cell Biology (203149) (to B.A.). We heartily thank Matúš Valach (University of Montreal, Montreal) for providing transcriptomes for 10 diplonemids. The authors would like to acknowledge the Proteomics Service Laboratory, Institute of Physiology and Institute of Molecular Genetics (Czech Academy of Sciences, Prague) and specifically Marek Vrbacký for mass spectrometry proteomics analysis, technical advice and expertise. We thank the members of the Kops group for detailed feedback. The authors declare no competing financial interests.

## Data availability

The mass spectrometry proteomics data have been deposited to the ProteomeXchange Consortium (http://proteomecentral.proteomexchange.org) via the PRIDE partner repository (108) with the dataset identifier PXD054535.

## Rights retention

This research was funded in whole, or in part, by the Wellcome Trust [227243/Z/23/Z]. For the purpose of open access, the author has applied a CC BY public copyright licence to any Author Accepted Manuscript version arising from this submission.

## REFERENCES

1. Chodasewicz K. 2014. Evolution, reproduction and definition of life. Theory Biosci 133:39–45.

2. Tetz VV, Tetz GV. 2020. A new biological definition of life. Biomol Concepts 11:1–6.

3. McIntosh JR. 2016. Mitosis. Cold Spring Harb Perspect Biol 8:a023218.

4. McAinsh AD, Kops GJPL. 2023. Principles and dynamics of spindle assembly checkpoint signalling. Nat Rev Mol Cell Biol 24:543–559.

5. Keeling PJ, Burki F. 2019. Progress towards the Tree of Eukaryotes. Curr Biol 29:R808–R817.

6. Drinnenberg IA, Akiyoshi B. 2017. Evolutionary lessons from species with unique kinetochores. Prog Mol Subcell Biol 56:111–138.

7. Musacchio A, Desai A. 2017. A molecular view of kinetochore assembly and function. Biology 6:5.

8. McAinsh AD, Marston AL. 2022. The four causes: The functional architecture of centromeres and kinetochores. Annu Rev Genet 56:279–314.

9. Meraldi P, McAinsh AD, Rheinbay E, Sorger PK. 2006. Phylogenetic and structural analysis of centromeric DNA and kinetochore proteins. Genome Biol 7:R23.

10. van Hooff JJ, Tromer E, van Wijk LM, Snel B, Kops GJ. 2017. Evolutionary dynamics of the kinetochore network in eukaryotes as revealed by comparative genomics. EMBO Rep 18:1559– 1571.

11. Tromer EC, van Hooff JJE, Kops GJPL, Snel B. 2019. Mosaic origin of the eukaryotic kinetochore. Proc Natl Acad Sci U S A 116:12873–12882.

12. Kops GJPL, Snel B, Tromer EC. 2020. Evolutionary dynamics of the spindle assembly checkpoint in eukaryotes. Curr Biol 30:R589–R602.

13. Ishii M, Akiyoshi B. 2022. Plasticity in centromere organization and kinetochore composition: Lessons from diversity. Curr Opin Cell Biol 74:47–54.

14. Drinnenberg IA, deYoung D, Henikoff S, Malik HS. 2014. Recurrent loss of CenH3 is associated with independent transitions to holocentricity in insects. eLife 3:e03676.

15. Navarro-Mendoza MI, Pérez-Arques C, Panchal S, Nicolás FE, Mondo SJ, Ganguly P, Pangilinan J, Grigoriev IV, Heitman J, Sanyal K, Garre V. 2019. Early diverging fungus *Mucor circinelloides* lacks centromeric histone CENP-A and displays a mosaic of point and regional centromeres. Curr Biol 29:3791–3802.e6.

16. Cortes-Silva N, Ulmer J, Kiuchi T, Hsieh E, Cornilleau G, Ladid I, Dingli F, Loew D, Katsuma S, Drinnenberg IA. 2020. CenH3-independent kinetochore assembly in Lepidoptera requires CCAN, including CENP-T. Curr Biol 30:561–572.e10.

17. Zeeshan M, Pandey R, Ferguson DJP, Tromer EC, Markus R, Abel S, Brady D, Daniel E, Limenitakis R, Bottrill AR, Le Roch KG, Holder AA, Waller RF, Guttery DS, Tewari R. 2020. Real-time dynamics of *Plasmodium* NDC80 reveals unusual modes of chromosome segregation during parasite proliferation. J Cell Sci 134:jcs245753.

18. Brusini L, Dos Santos Pacheco N, Tromer EC, Soldati-Favre D, Brochet M. 2022. Composition and organization of kinetochores show plasticity in apicomplexan chromosome segregation. J Cell Biol 221:e202111084.

19. Salas-Leiva DE, Tromer EC, Curtis BA, Jerlström-Hultqvist J, Kolisko M, Yi Z, Salas-Leiva JS, Gallot-Lavallée L, Williams SK, Kops GJPL, Archibald JM, Simpson AGB, Roger AJ. 2021. Genomic analysis finds no evidence of canonical eukaryotic DNA processing complexes in a free-living protist. Nat Commun 12:6003.

20. Berriman M, Ghedin E, Hertz-Fowler C, Blandin G, Renauld H, Bartholomeu DC, Lennard NJ, Caler E, Hamlin NE, Haas B, Böhme U, Hannick L, Aslett MA, Shallom J, Marcello L, Hou L, Wickstead B, Alsmark UCM, Arrowsmith C, Atkin RJ, Barron AJ, Bringaud F, Brooks K, Carrington M, Cherevach I, Chillingworth T-J, Churcher C, Clark LN, Corton CH, Cronin A, Davies RM, Doggett J, Djikeng A, Feldblyum T, Field MC, Fraser A, Goodhead I, Hance Z, Harper D, Harris BR, Hauser H, Hostetler J, Ivens A, Jagels K, Johnson D, Johnson J, Jones K, Kerhornou AX, Koo H, Larke N, Landfear S, Larkin C, Leech V, Line A, Lord A, Macleod A, Mooney PJ, Moule S, Martin DMA, Morgan GW, Mungall K, Norbertczak H, Ormond D, Pai G, Peacock CS, Peterson J, Quail MA, Rabbinowitsch E, Rajandream M-A, Reitter C, Salzberg SL, Sanders M, Schobel S, Sharp S, Simmonds M, Simpson AJ, Tallon L, Turner CMR, Tait A, Tivey AR, Van Aken S, Walker D, Wanless D, Wang S, White B, White O, Whitehead S, Woodward J, Wortman J, Adams MD, Embley TM, Gull K, Ullu E, Barry JD, Fairlamb AH, Opperdoes F, Barrell BG, Donelson JE, Hall N, Fraser CM, Melville SE, El-Sayed NM. 2005. The genome of the African trypanosome *Trypanosoma brucei*. Science 309:416–422.

21. Akiyoshi B, Gull K. 2013. Evolutionary cell biology of chromosome segregation: insights from trypanosomes. Open Biol 3:130023.

22. Akiyoshi B, Gull K. 2014. Discovery of unconventional kinetochores in kinetoplastids. Cell 156:1247–1258.

23. Nerusheva OO, Akiyoshi B. 2016. Divergent polo box domains underpin the unique kinetoplastid kinetochore. Open Biol 6:150206.

24. D’Archivio S, Wickstead B. 2017. Trypanosome outer kinetochore proteins suggest conservation of chromosome segregation machinery across eukaryotes. J Cell Biol 216:379– 391.

25. Nerusheva OO, Ludzia P, Akiyoshi B. 2019. Identification of four unconventional kinetoplastid kinetochore proteins KKT22-25 in *Trypanosoma brucei*. Open Biol 9:190236.

26. Brusini L, D’Archivio S, McDonald J, Wickstead B. 2021. Trypanosome KKIP1 dynamically links the inner kinetochore to a kinetoplastid outer kinetochore complex. Front Cell Infect Microbiol 11:641174.

27. Aslett M, Aurrecoechea C, Berriman M, Brestelli J, Brunk BP, Carrington M, Depledge DP, Fischer S, Gajria B, Gao X, Gardner MJ, Gingle A, Grant G, Harb OS, Heiges M, Hertz-Fowler C, Houston R, Innamorato F, Iodice J, Kissinger JC, Kraemer E, Li W, Logan FJ, Miller JA, Mitra S, Myler PJ, Nayak V, Pennington C, Phan I, Pinney DF, Ramasamy G, Rogers MB, Roos DS, Ross C, Sivam D, Smith DF, Srinivasamoorthy G, Stoeckert CJ, Subramanian S, Thibodeau R, Tivey A, Treatman C, Velarde G, Wang H. 2010. TriTrypDB: a functional genomic resource for the Trypanosomatidae. Nucleic Acids Res 38:D457–462.

28. Jackson AP, Otto TD, Aslett M, Armstrong SD, Bringaud F, Schlacht A, Hartley C, Sanders M, Wastling JM, Dacks JB, Acosta-Serrano A, Field MC, Ginger ML, Berriman M. 2016. Kinetoplastid phylogenomics reveals the evolutionary innovations associated with the origins of parasitism. Curr Biol 26:161–172.

29. Butenko A, Opperdoes FR, Flegontova O, Horák A, Hampl V, Keeling P, Gawryluk RMR, Tikhonenkov D, Flegontov P, Lukeš J. 2020. Evolution of metabolic capabilities and molecular features of diplonemids, kinetoplastids, and euglenids. BMC Biol 18:23.

30. Tikhonenkov DV, Gawryluk RMR, Mylnikov AP, Keeling PJ. 2021. First finding of free-living representatives of Prokinetoplastina and their nuclear and mitochondrial genomes. Sci Rep 11:2946.

31. Tromer EC, Wemyss TA, Ludzia P, Waller RF, Akiyoshi B. 2021. Repurposing of synaptonemal complex proteins for kinetochores in Kinetoplastida. Open Biol 11:210049.

32. Cavalier-Smith T. 2016. Higher classification and phylogeny of Euglenozoa. Eur J Protistol 56:250–276.

33. Kostygov AY, Karnkowska A, Votýpka J, Tashyreva D, Maciszewski K, Yurchenko V, Lukeš J. 2021. Euglenozoa: taxonomy, diversity and ecology, symbioses and viruses. Open Biol 11:200407.

34. Lax G, Kolisko M, Eglit Y, Lee WJ, Yubuki N, Karnkowska A, Leander BS, Burger G, Keeling PJ, Simpson AGB. 2021. Multigene phylogenetics of euglenids based on single-cell transcriptomics of diverse phagotrophs. Mol Phylogenet Evol 159:107088.

35. de Vargas C, Audic S, Henry N, Decelle J, Mahé F, Logares R, Lara E, Berney C, Le Bescot N, Probert I, Carmichael M, Poulain J, Romac S, Colin S, Aury J-M, Bittner L, Chaffron S, Dunthorn M, Engelen S, Flegontova O, Guidi L, Horák A, Jaillon O, Lima-Mendez G, Lukeš J, Malviya S, Morard R, Mulot M, Scalco E, Siano R, Vincent F, Zingone A, Dimier C, Picheral M, Searson S, Kandels-Lewis S, Tara Oceans Coordinators, Acinas SG, Bork P, Bowler C, Gorsky G, Grimsley N, Hingamp P, Iudicone D, Not F, Ogata H, Pesant S, Raes J, Sieracki ME, Speich S, Stemmann L, Sunagawa S, Weissenbach J, Wincker P, Karsenti E. 2015. Ocean plankton. Eukaryotic plankton diversity in the sunlit ocean. Science 348:1261605.

36. Flegontova O, Flegontov P, Malviya S, Audic S, Wincker P, de Vargas C, Bowler C, Lukeš J, Horák A. 2016. Extreme diversity of Diplonemid eukaryotes in the ocean. Curr Biol 26:3060– 3065.

37. Tashyreva D, Simpson AGB, Prokopchuk G, Škodová-Sveráková I, Butenko A, Hammond M, George EE, Flegontova O, Záhonová K, Faktorová D, Yabuki A, Horák A, Keeling PJ, Lukeš J. 2022. Diplonemids - A review on “new” flagellates on the oceanic block. Protist 173:125868.

38. Flegontova O, Flegontov P, Londoño PAC, Walczowski W, Šantić D, Edgcomb VP, Lukeš J, Horák A. 2020. Environmental determinants of the distribution of planktonic diplonemids and kinetoplastids in the oceans. Environ Microbiol 22:4014–4031.

39. Tashyreva D, Týč J, Horák A, Lukeš J. 2023. Ultrastructure and 3D reconstruction of a diplonemid protist (Diplonemea) and its novel membranous organelle. mBio 14:e0192123.

40. van Hooff JJE, Raas MWD, Tromer E, Eme L. 2024. Shaping up genomes: Prokaryotic roots and eukaryotic diversification of SMC complexes. bioRxiv 10.1101/2024.01.07.573240.

41. Kaur B, Valach M, Peña-Diaz P, Moreira S, Keeling PJ, Burger G, Lukeš J, Faktorová D. 2018. Transformation of *Diplonema papillatum*, the type species of the highly diverse and abundant marine microeukaryotes Diplonemida (Euglenozoa). Environ Microbiol 20:1030–1040.

42. Faktorová D, Nisbet RER, Fernández Robledo JA, Casacuberta E, Sudek L, Allen AE, Ares M, Aresté C, Balestreri C, Barbrook AC, Beardslee P, Bender S, Booth DS, Bouget F-Y, Bowler C, Breglia SA, Brownlee C, Burger G, Cerutti H, Cesaroni R, Chiurillo MA, Clemente T, Coles DB, Collier JL, Cooney EC, Coyne K, Docampo R, Dupont CL, Edgcomb V, Einarsson E, Elustondo PA, Federici F, Freire-Beneitez V, Freyria NJ, Fukuda K, García PA, Girguis PR, Gomaa F, Gornik SG, Guo J, Hampl V, Hanawa Y, Haro-Contreras ER, Hehenberger E, Highfield A, Hirakawa Y, Hopes A, Howe CJ, Hu I, Ibañez J, Irwin NAT, Ishii Y, Janowicz NE, Jones AC, Kachale A, Fujimura-Kamada K, Kaur B, Kaye JZ, Kazana E, Keeling PJ, King N, Klobutcher LA, Lander N, Lassadi I, Li Z, Lin S, Lozano J-C, Luan F, Maruyama S, Matute T, Miceli C, Minagawa J, Moosburner M, Najle SR, Nanjappa D, Nimmo IC, Noble L, Novák Vanclová AMG, Nowacki M, Nuñez I, Pain A, Piersanti A, Pucciarelli S, Pyrih J, Rest JS, Rius M, Robertson D, Ruaud A, Ruiz-Trillo I, Sigg MA, Silver PA, Slamovits CH, Jason Smith G, Sprecher BN, Stern R, Swart EC, Tsaousis AD, Tsypin L, Turkewitz A, Turnšek J, Valach M, Vergé V, von Dassow P, von der Haar T, Waller RF, Wang L, Wen X, Wheeler G, Woods A, Zhang H, Mock T, Worden AZ, Lukeš J. 2020. Genetic tool development in marine protists: emerging model organisms for experimental cell biology. Nat Methods 17:481–494.

43. Faktorová D, Kaur B, Valach M, Graf L, Benz C, Burger G, Lukeš J. 2020. Targeted integration by homologous recombination enables in situ tagging and replacement of genes in the marine microeukaryote *Diplonema papillatum*. Environ Microbiol 22:3660–3670.

44. Faktorová D, Záhonová K, Benz C, Dacks JB, Field MC, Lukeš J. 2023. Functional differentiation of Sec13 paralogues in the euglenozoan protists. Open Biol 13:220364.

45. Valach M, Benz C, Aguilar LC, Gahura O, Faktorová D, Zíková A, Oeffinger M, Burger G, Gray MW, Lukeš J. 2023. Miniature RNAs are embedded in an exceptionally protein-rich mitoribosome via an elaborate assembly pathway. Nucleic Acids Res 51:6443–6460.

46. Valach M, Moreira S, Petitjean C, Benz C, Butenko A, Flegontova O, Nenarokova A, Prokopchuk G, Batstone T, Lapébie P, Lemogo L, Sarrasin M, Stretenowich P, Tripathi P, Yazaki E, Nara T, Henrissat B, Lang BF, Gray MW, Williams TA, Lukeš J, Burger G. 2023. Recent expansion of metabolic versatility in *Diplonema papillatum*, the model species of a highly speciose group of marine eukaryotes. BMC Biol 21:99.

47. Škodová-Sveráková I, Záhonová K, Juricová V, Danchenko M, Moos M, Baráth P, Prokopchuk G, Butenko A, Lukáčová V, Kohútová L, Bučková B, Horák A, Faktorová D, Horváth A, Šimek P, Lukeš J. 2021. Highly flexible metabolism of the marine euglenozoan protist *Diplonema papillatum*. BMC Biol 19:251.

48. Wells JN, Marsh JA. 2019. A graph-based approach for detecting sequence homology in highly diverged repeat protein families, p. 251–261. In Sikosek, T (ed.), Computational Methods in Protein Evolution. Springer, New York, NY.

49. Schou KB, Andersen JS, Pedersen LB. 2014. A divergent calponin homology (NN-CH) domain defines a novel family: implications for evolution of ciliary IFT complex B proteins. Bioinformatics 30:899–902.

50. Holden JM, Koreny L, Obado S, Ratushny AV, Chen W-M, Chiang J-H, Kelly S, Chait BT, Aitchison JD, Rout MP, Field MC. 2014. Nuclear pore complex evolution: a trypanosome Mlp analogue functions in chromosomal segregation but lacks transcriptional barrier activity. Mol Biol Cell 25:1421–1436.

51. Ballmer D, Carter W, van Hooff JJE, Tromer EC, Ishii M, Ludzia P, Akiyoshi B. 2024. Kinetoplastid kinetochore proteins KKT14-KKT15 are divergent Bub1/BubR1-Bub3 proteins. Open Biol 14:240025.

52. Wijk LM van, Snel B. 2020. The first eukaryotic kinome tree illuminates the dynamic history of present-day kinases. bioRxiv 10.1101/2020.01.27.920793.

53. Derelle R, Torruella G, Klimeš V, Brinkmann H, Kim E, Vlček Č, Lang BF, Eliáš M. 2015. Bacterial proteins pinpoint a single eukaryotic root. Proc Natl Acad Sci U S A 112:E693–699.

54. Stok C, Tsaridou S, van den Tempel N, Everts M, Wierenga E, Bakker FJ, Kok Y, Alves IT, Jae LT, Raas MWD, Huis In’t Veld PJ, de Boer HR, Bhattacharya A, Karanika E, Warner H, Chen M, van de Kooij B, Dessapt J, Ter Morsche L, Perepelkina P, Fradet-Turcotte A, Guryev V, Tromer EC, Chan K-L, Fehrmann RSN, van Vugt MATM. 2023. FIRRM/C1orf112 is synthetic lethal with PICH and mediates RAD51 dynamics. Cell Rep 42:112668.

55. van Rooijen LE, Tromer EC, van Hooff JJE, Kops GJPL, Snel B. 2023. Increased sampling and intracomplex homologies favor vertical over horizontal inheritance of the Dam1 complex. Genome Biol Evol 15:evad017.

56. Burki F, Roger AJ, Brown MW, Simpson AGB. 2020. The new tree of eukaryotes. Trends Ecol Evol 35:43–55.

57. Mansfeld J, Collin P, Collins MO, Choudhary JS, Pines J. 2011. APC15 drives the turnover of MCC-CDC20 to make the spindle assembly checkpoint responsive to kinetochore attachment. Nat Cell Biol 13:1234–1243.

58. Osman F, Whitby MC. 2013. Emerging roles for centromere-associated proteins in DNA repair and genetic recombination. Biochem Soc Trans 41:1726–1730.

59. Malik HS, Henikoff S. 2003. Phylogenomics of the nucleosome. Nat Struct Biol 10:882–891.

60. Luo X, Tang Z, Rizo J, Yu H. 2002. The Mad2 spindle checkpoint protein undergoes similar major conformational changes upon binding to either Mad1 or Cdc20. Mol Cell 9:59–71.

61. Aravind L, Koonin EV. 1998. The HORMA domain: a common structural denominator in mitotic checkpoints, chromosome synapsis and DNA repair. Trends Biochem Sci 23:284–286.

62. Carmena M, Wheelock M, Funabiki H, Earnshaw WC. 2012. The chromosomal passenger complex (CPC): from easy rider to the godfather of mitosis. Nat Rev Mol Cell Biol 13:789– 803.

63. Li Z, Lee JH, Chu F, Burlingame AL, Günzl A, Wang CC. 2008. Identification of a novel chromosomal passenger complex and its unique localization during cytokinesis in *Trypanosoma brucei*. PLoS One 3:e2354.

64. Ballmer D, Akiyoshi B. 2024. Dynamic localization of the chromosomal passenger complex in trypanosomes is controlled by the orphan kinesins KIN-A and KIN-B. eLife 13:RP93522.

65. Adams RR, Wheatley SP, Gouldsworthy AM, Kandels-Lewis SE, Carmena M, Smythe C, Gerloff DL, Earnshaw WC. 2000. INCENP binds the Aurora-related kinase AIRK2 and is required to target it to chromosomes, the central spindle and cleavage furrow. Curr Biol 10:1075–1078.

66. Sessa F, Mapelli M, Ciferri C, Tarricone C, Areces LB, Schneider TR, Stukenberg PT, Musacchio A. 2005. Mechanism of Aurora B activation by INCENP and inhibition by hesperadin. Mol Cell 18:379–391.

67. Abramson J, Adler J, Dunger J, Evans R, Green T, Pritzel A, Ronneberger O, Willmore L, Ballard AJ, Bambrick J, Bodenstein SW, Evans DA, Hung C-C, O’Neill M, Reiman D, Tunyasuvunakool K, Wu Z, Žemgulytė A, Arvaniti E, Beattie C, Bertolli O, Bridgland A, Cherepanov A, Congreve M, Cowen-Rivers AI, Cowie A, Figurnov M, Fuchs FB, Gladman H, Jain R, Khan YA, Low CMR, Perlin K, Potapenko A, Savy P, Singh S, Stecula A, Thillaisundaram A, Tong C, Yakneen S, Zhong ED, Zielinski M, Žídek A, Bapst V, Kohli P, Jaderberg M, Hassabis D, Jumper JM. 2024. Accurate structure prediction of biomolecular interactions with AlphaFold 3. Nature 630:493–500.

68. Ochs RL, Press RI. 1992. Centromere autoantigens are associated with the nucleolus. Exp Cell Res 200:339–350.

69. Bury L, Moodie B, Ly J, McKay LS, Miga KH, Cheeseman IM. 2020. Alpha-satellite RNA transcripts are repressed by centromere-nucleolus associations. eLife 9:e59770.

70. Rodrigues A, MacQuarrie KL, Freeman E, Lin A, Willis AB, Xu Z, Alvarez AA, Ma Y, White BEP, Foltz DR, Huang S. 2023. Nucleoli and the nucleoli-centromere association are dynamic during normal development and in cancer. Mol Biol Cell 34:br5.

71. Komaki S, Tromer EC, De Jaeger G, De Winne N, Heese M, Schnittger A. 2022. Molecular convergence by differential domain acquisition is a hallmark of chromosomal passenger complex evolution. Proc Natl Acad Sci U S A 119:e2200108119.

72. Zickler D, Kleckner N. 2023. Meiosis: Dances between homologs. Annu Rev Genet 57:1–63.

73. Ishii M, Akiyoshi B. 2020. Characterization of unconventional kinetochore kinases KKT10 and KKT19 in *Trypanosoma brucei*. J Cell Sci 133:jcs240978.

74. Hardie DG. 1999. Plant protein serine/threonine kinases: Classification and functions. Annu Rev Plant Physiol Plant Mol Biol 50:97–131.

75. Lindberg MF, Meijer L. 2021. Dual-specificity, tyrosine phosphorylation-regulated kinases (DYRKs) and cdc2-like kinases (CLKs) in human disease, an overview. Int J Mol Sci 22:6047.

76. Garcia-Silva M-R, Sollelis L, MacPherson CR, Stanojcic S, Kuk N, Crobu L, Bringaud F, Bastien P, Pagès M, Scherf A, Sterkers Y. 2017. Identification of the centromeres of *Leishmania major*: revealing the hidden pieces. EMBO Rep 18:1968–1977.

77. Geoghegan V, Carnielli JBT, Jones NG, Saldivia M, Antoniou S, Hughes C, Neish R, Dowle A, Mottram JC. 2022. CLK1/CLK2-driven signalling at the *Leishmania* kinetochore is captured by spatially referenced proximity phosphoproteomics. Commun Biol 5:1305.

78. Porter D. 1973. Isonema papillatum sp. n., a new colorless marine flagellate: A light- and electronmicroscopic study. J Protozool 20:351–356.

79. Triemer RE. 1992. Ultrastructure of mitosis in *Diplonema ambulator* Larsen and Patterson (Euglenozoa). Eur J Protistol 28:398–404.

80. Lowell JE, Cross GAM. 2004. A variant histone H3 is enriched at telomeres in *Trypanosoma brucei*. J Cell Sci 117:5937–5947.

81. Alfieri C, Chang L, Barford D. 2018. Mechanism for remodelling of the cell cycle checkpoint protein MAD2 by the ATPase TRIP13. Nature 559:274–278.

82. Butterfield ER, Obado SO, Scutts SR, Zhang W, Chait BT, Rout MP, Field MC. 2024. A lineage-specific protein network at the trypanosome nuclear envelope. Nucleus 15:2310452.

83. Ebenezer TE, Zoltner M, Burrell A, Nenarokova A, Novák Vanclová AMG, Prasad B, Soukal P, Santana-Molina C, O’Neill E, Nankissoor NN, Vadakedath N, Daiker V, Obado S, Silva-Pereira S, Jackson AP, Devos DP, Lukeš J, Lebert M, Vaughan S, Hampl V, Carrington M, Ginger ML, Dacks JB, Kelly S, Field MC. 2019. Transcriptome, proteome and draft genome of *Euglena gracilis*. BMC Biol 17:11.

84. Pastor F, Shkreta L, Chabot B, Durantel D, Salvetti A. 2021. Interplay between CMGC kinases targeting SR proteins and viral replication: Splicing and beyond. Front Microbiol 12:658721.

85. Lukeš J, Speijer D, Zíková A, Alfonzo JD, Hashimi H, Field MC. 2023. Trypanosomes as a magnifying glass for cell and molecular biology. Trends Parasitol 39:902–912.

86. Kaur B, Záhonová K, Valach M, Faktorová D, Prokopchuk G, Burger G, Lukeš J. 2020. Gene fragmentation and RNA editing without borders: eccentric mitochondrial genomes of diplonemids. Nucleic Acids Res 48:2694–2708.

87. Valach M, Moreira S, Hoffmann S, Stadler PF, Burger G. 2017. Keeping it complicated: Mitochondrial genome plasticity across diplonemids. Sci Rep 7:14166.

88. George EE, Tashyreva D, Kwong WK, Okamoto N, Horák A, Husnik F, Lukeš J, Keeling PJ. 2022. Gene transfer agents in bacterial endosymbionts of microbial eukaryotes. Genome Biol Evol 14:evac099.

89. Haas BJ, Papanicolaou A, Yassour M, Grabherr M, Blood PD, Bowden J, Couger MB, Eccles D, Li B, Lieber M, MacManes MD, Ott M, Orvis J, Pochet N, Strozzi F, Weeks N, Westerman R, William T, Dewey CN, Henschel R, LeDuc RD, Friedman N, Regev A. 2013. De novo transcript sequence reconstruction from RNA-seq using the Trinity platform for reference generation and analysis. Nat Protoc 8:1494–1512.

90. Huang Y, Niu B, Gao Y, Fu L, Li W. 2010. CD-HIT Suite: a web server for clustering and comparing biological sequences. Bioinformatics 26:680–682.

91. Jumper J, Evans R, Pritzel A, Green T, Figurnov M, Ronneberger O, Tunyasuvunakool K, Bates R, Žídek A, Potapenko A, Bridgland A, Meyer C, Kohl SAA, Ballard AJ, Cowie A, Romera-Paredes B, Nikolov S, Jain R, Adler J, Back T, Petersen S, Reiman D, Clancy E, Zielinski M, Steinegger M, Pacholska M, Berghammer T, Bodenstein S, Silver D, Vinyals O, Senior AW, Kavukcuoglu K, Kohli P, Hassabis D. 2021. Highly accurate protein structure prediction with AlphaFold. Nature 596:583–589.

92. Wheeler RJ. 2021. A resource for improved predictions of *Trypanosoma* and *Leishmania* protein three-dimensional structure. PLoS One 16:e0259871.

93. Steinegger M, Söding J. 2017. MMseqs2 enables sensitive protein sequence searching for the analysis of massive data sets. Nat Biotechnol 35:1026–1028.

94. Richter DJ, Berney C, Strassert JFH, Poh Y-P, Herman EK, Muñoz-Gómez SA, Wideman JG, Burki F, de Vargas C. 2022. EukProt: A database of genome-scale predicted proteins across the diversity of eukaryotes. Peer J 2:e56.

95. Mirdita M, Schütze K, Moriwaki Y, Heo L, Ovchinnikov S, Steinegger M. 2022. ColabFold: making protein folding accessible to all. Nat Methods 19:679–682.

96. Eddy SR. 1998. Profile hidden Markov models. Bioinformatics 14:755–763.

97. Emms DM, Kelly S. 2019. OrthoFinder: phylogenetic orthology inference for comparative genomics. Genome Biol 20:238.

98. Katoh K, Standley DM. 2013. MAFFT multiple sequence alignment software version 7: improvements in performance and usability. Mol Biol Evol 30:772–780.

99. Steinegger M, Meier M, Mirdita M, Vöhringer H, Haunsberger SJ, Söding J. 2019. HH-suite3 for fast remote homology detection and deep protein annotation. BMC Bioinformatics 20:473.

100. Shannon P, Markiel A, Ozier O, Baliga NS, Wang JT, Ramage D, Amin N, Schwikowski B, Ideker T. 2003. Cytoscape: a software environment for integrated models of biomolecular interaction networks. Genome Res 13:2498–2504.

101. van Kempen M, Kim SS, Tumescheit C, Mirdita M, Lee J, Gilchrist CLM, Söding J, Steinegger M. 2024. Fast and accurate protein structure search with Foldseek. Nat Biotechnol 42:243–246.

102. de Potter B, Raas MWD, Seidl MF, Verrijzer CP, Snel B. 2023. Uncoupled evolution of the Polycomb system and deep origin of non-canonical PRC1. Commun Biol 6:1144.

103. Minh BQ, Schmidt HA, Chernomor O, Schrempf D, Woodhams MD, von Haeseler A, Lanfear R. 2020. IQ-TREE 2: New models and efficient methods for phylogenetic inference in the genomic era. Mol Biol Evol 37:1530–1534.

104. Kalyaanamoorthy S, Minh BQ, Wong TKF, von Haeseler A, Jermiin LS. 2017. ModelFinder: fast model selection for accurate phylogenetic estimates. Nat Methods 14:587–589.

105. Masuda T, Tomita M, Ishihama Y. 2008. Phase transfer surfactant-aided trypsin digestion for membrane proteome analysis. J Proteome Res 7:731–740.

106. Cox J, Mann M. 2008. MaxQuant enables high peptide identification rates, individualized p.p.b.-range mass accuracies and proteome-wide protein quantification. Nat Biotechnol 26:1367–1372.

107. Tyanova S, Cox J. 2018. Perseus: A Bioinformatics Platform for Integrative Analysis of Proteomics Data in Cancer Research, p. 133–148. *In* von Stechow, L (ed.), Cancer Systems Biology: Methods and Protocols. Springer, New York, NY.

108. Perez-Riverol Y, Bai J, Bandla C, García-Seisdedos D, Hewapathirana S, Kamatchinathan S, Kundu DJ, Prakash A, Frericks-Zipper A, Eisenacher M, Walzer M, Wang S, Brazma A, Vizcaíno JA. 2022. The PRIDE database resources in 2022: a hub for mass spectrometry-based proteomics evidences. Nucleic Acids Res 50:D543–D552.

