## Supplementary material for "On the possibility of yet a third kinetochore system in the protist phylum Euglenozoa"

#### FIGURES

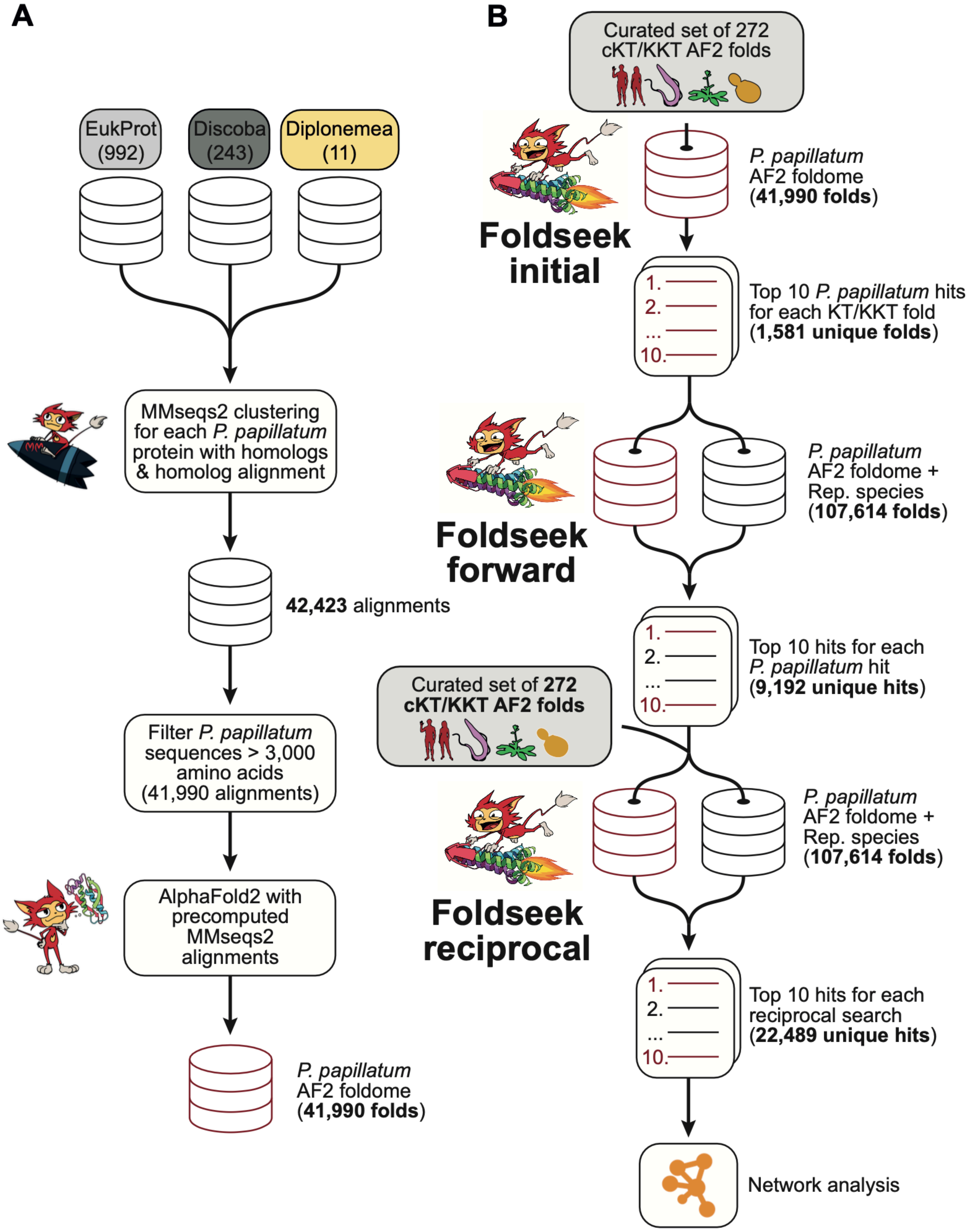


**Fig. S1 Bioinformatic pipeline overview of structure-based homology searching.** (A) Approach to produce high-quality structure predictions for all *P. papillatum* accessions using AlphaFold2 using custom alignments enriched for discoban & diplonemid homologs. (B) Strategy for reciprocal homology searching using Foldseek towards identifying divergent *c*KT/KKT orthologs.

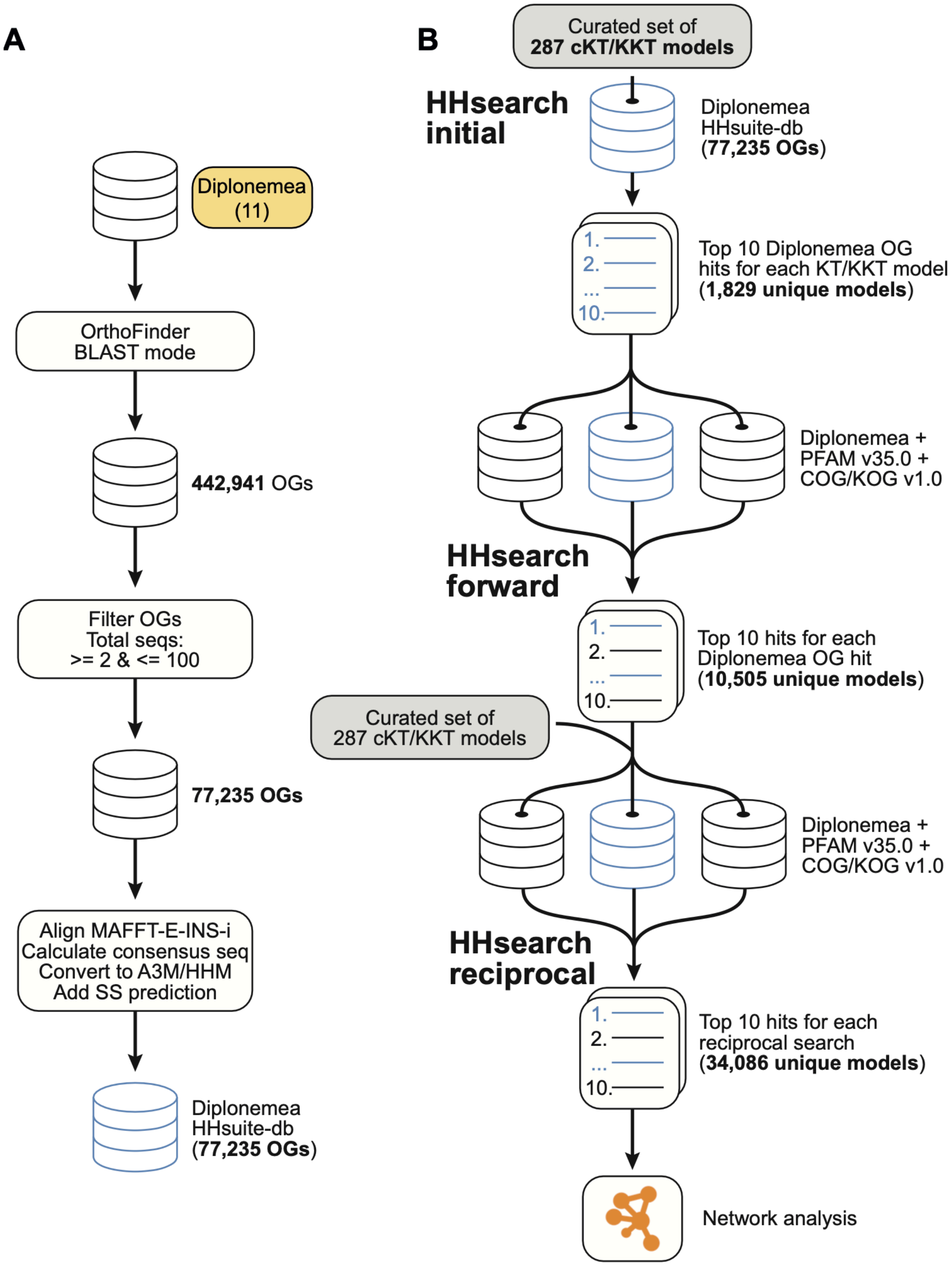


**Fig. S2 Bioinformatic pipeline overview of HMM-vs-HMM-based homology searching.** (A) Approach to produce a comprehensive alignment database of orthologous groups from diplonemid predicted proteomes. (B) Strategy for reciprocal homology searching using HHsearch and HMM-vs-HMM searching towards identifying divergent *c*KT/KKT orthologs.

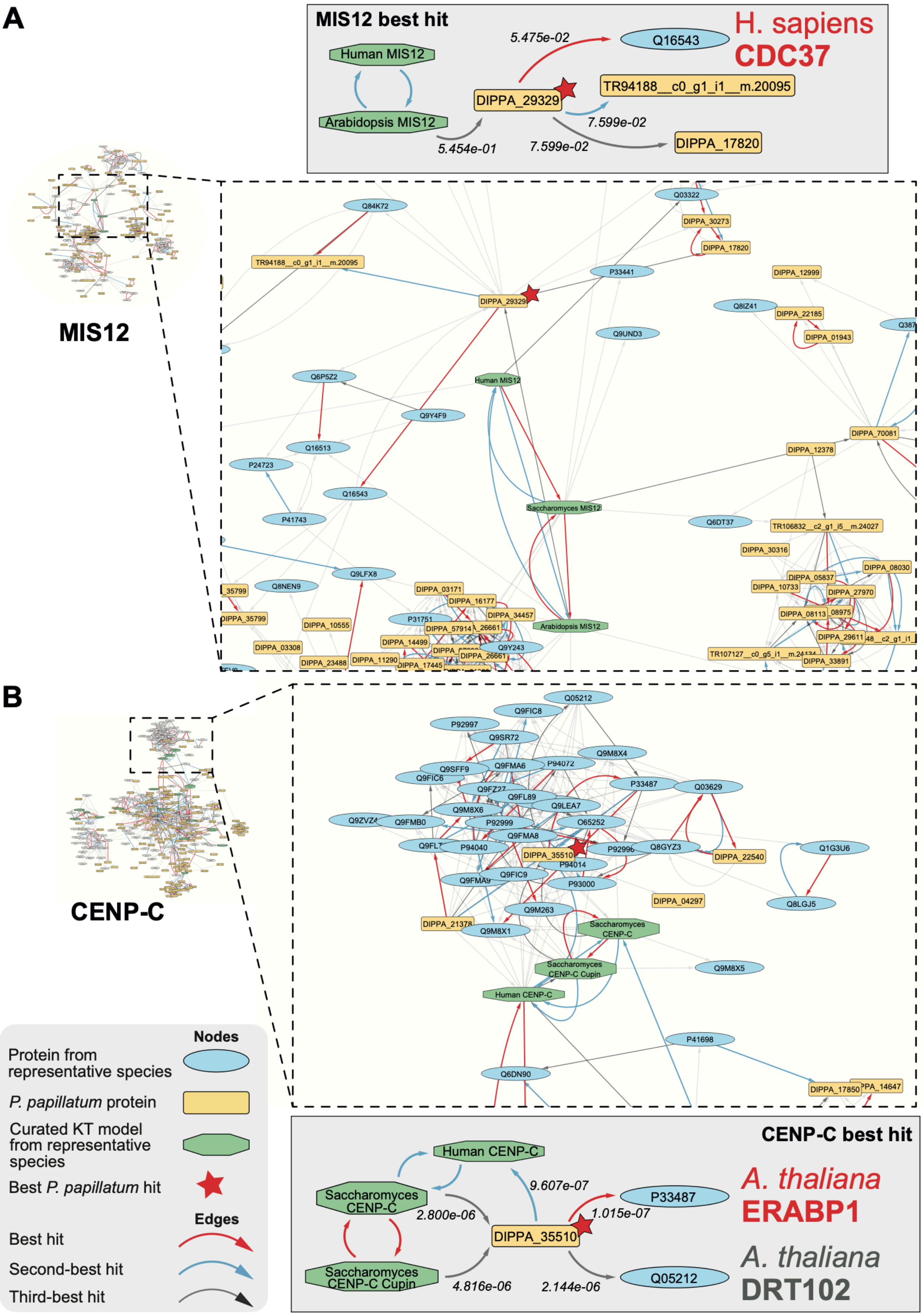


**Fig. S3 Best hits to *c*KT CENP-C and MIS12 in *P. papillatum* are not 1-to-1 orthologs but other such proteins with similar folds.** (A-B) Network graphs showing homology hits as inferred by

Foldseek, displayed as edges between homologous folds depicted as nodes. (A) All nodes connected to MIS12 nodes through either incoming or outgoing edges, and all nodes connected to these nodes with either directionality to the edge (i.e. all nodes until one degree of separation to a MIS12 node). Nodes are positioned relative to other nodes according to the E-value represented by connecting edges, where shorter edges equate to lower E-value. On the left, a zoom-in view of the network around the node of the *P. papillatum* accession that is best hit by one of the MIS12 query models (marked by a red star) and its most closely-related hits. Schematic representations of the path connecting MIS12 query nodes to the top *P. papillatum* accession hit and its subsequent best hits are shown. (B) Similar to panel A, but with nodes connected to CENP-C through maximally one degree of separation.

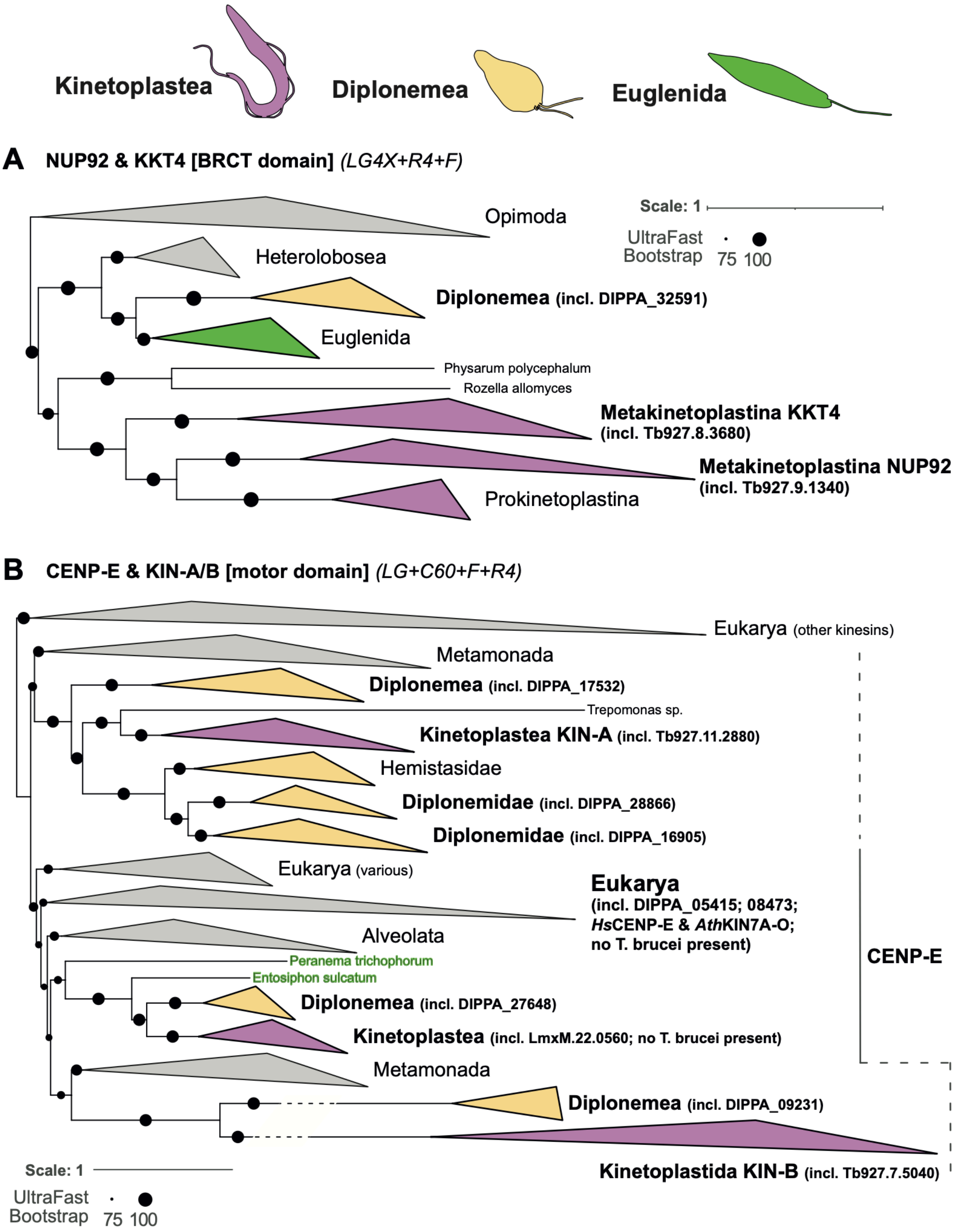


**Fig. S4 Phylogenetic trees of NUP92/KKT4 and CENP-E/KIN-A/B.** (A) Phylogram depicting the inferred phylogeny between NUP92 and KKT4 among Euglenozoa, including other eukaryotic outgroup proteins (NUP92). The tree was modelled using the LG4X+R4+F substitution model. (B) Phylogenetic tree depicting the inferred phylogeny of CENP-E/KIN-A/B motor domains and homologous sequences as identified in an extensive set of eukaryote-wide proteomes, including an extended set of Discoba proteomes. The tree was modelled using the LG+C60+F+R4 substitution model.

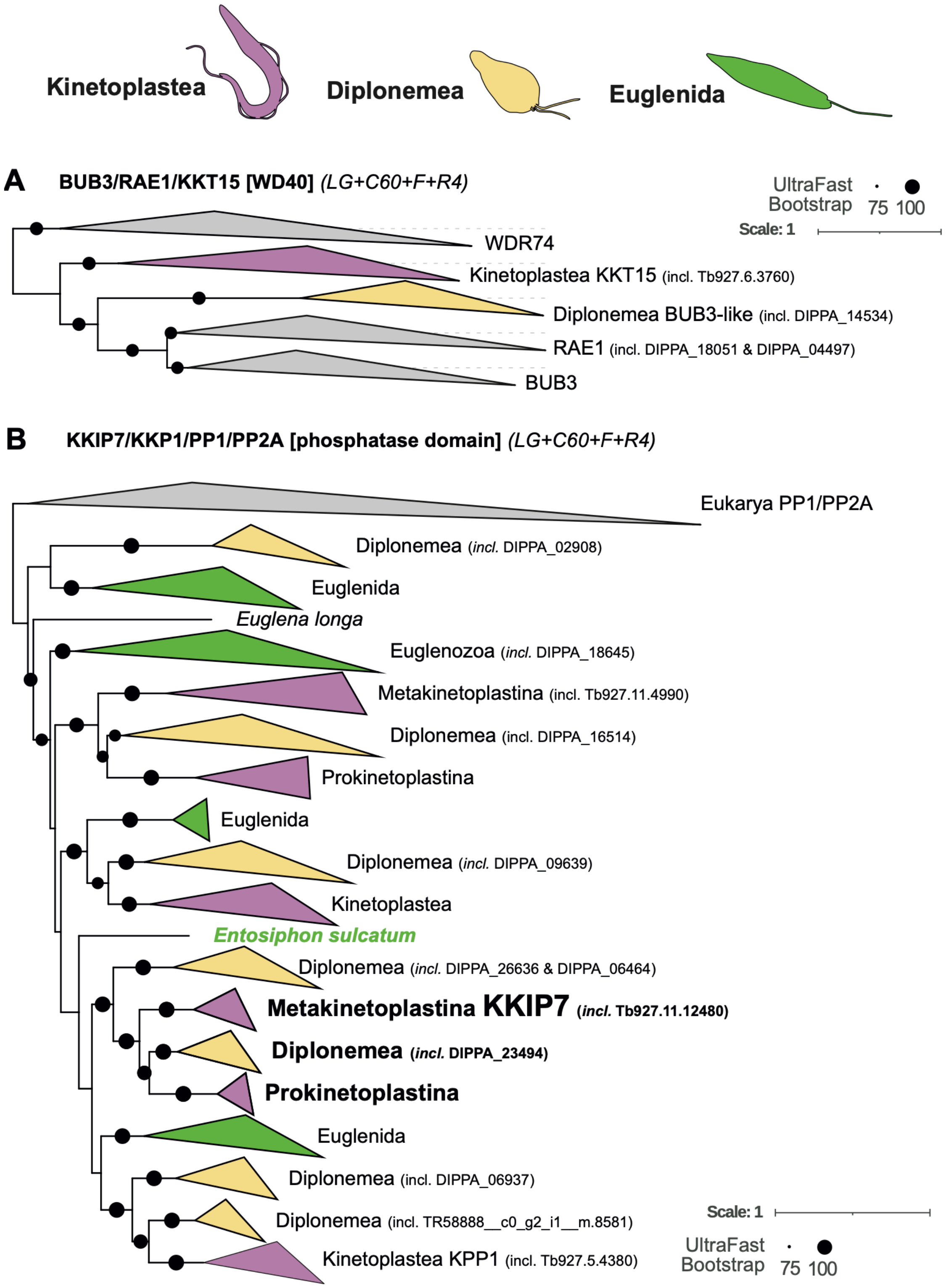


**Fig. S5 Phylogenetic analysis of BUB3 & KKIP7 homologs.** (A) Phylogenetic tree depicting the evolutionary relationships between BUB3, RAE1, KKT15, WDR74 and a putative BUB3-like set of accessions from Diplonemea. The tree was modelled using the LG+C60+F+R4 substitution model. (B) Phylogenetic tree depicting the inferred phylogeny of KKIP7 and homologous sequences as identified in an extensive set of eukaryote-wide proteomes, including an extended set of Discoba proteomes. The tree was modelled using the LG+C60+F+R4 substitution model.

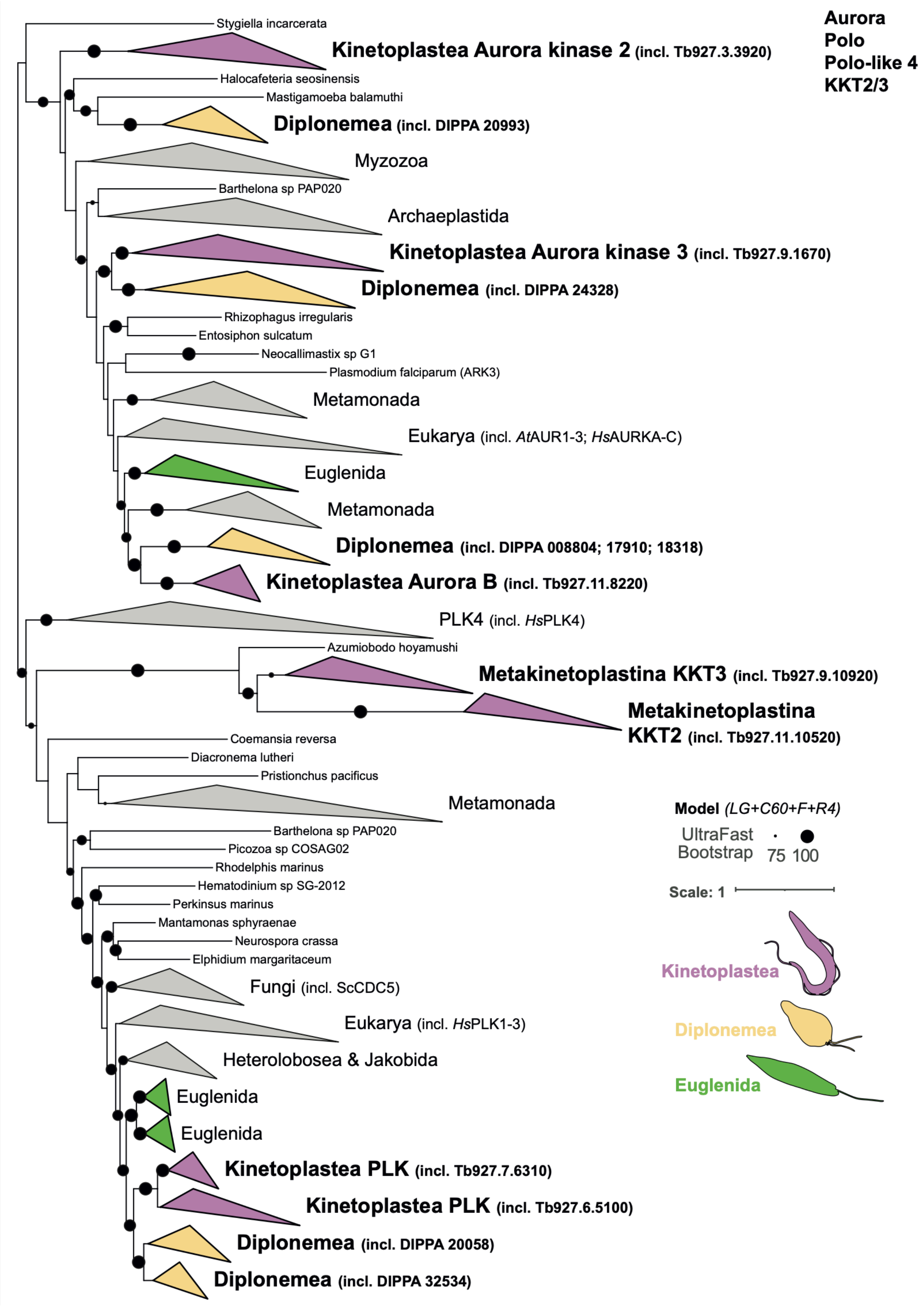


**Fig. S6 Phylogenetic tree of Aurora/Polo kinase.** Phylogram depicting the inferred kinase domain phylogeny of Aurora, Polo, PLK4, KKT2 and KKT3 for Discoba, supplemented with diverse sequences from various eukaryotic clades (11). The tree was modelled using the LG+C60+F+R4 substitution model.

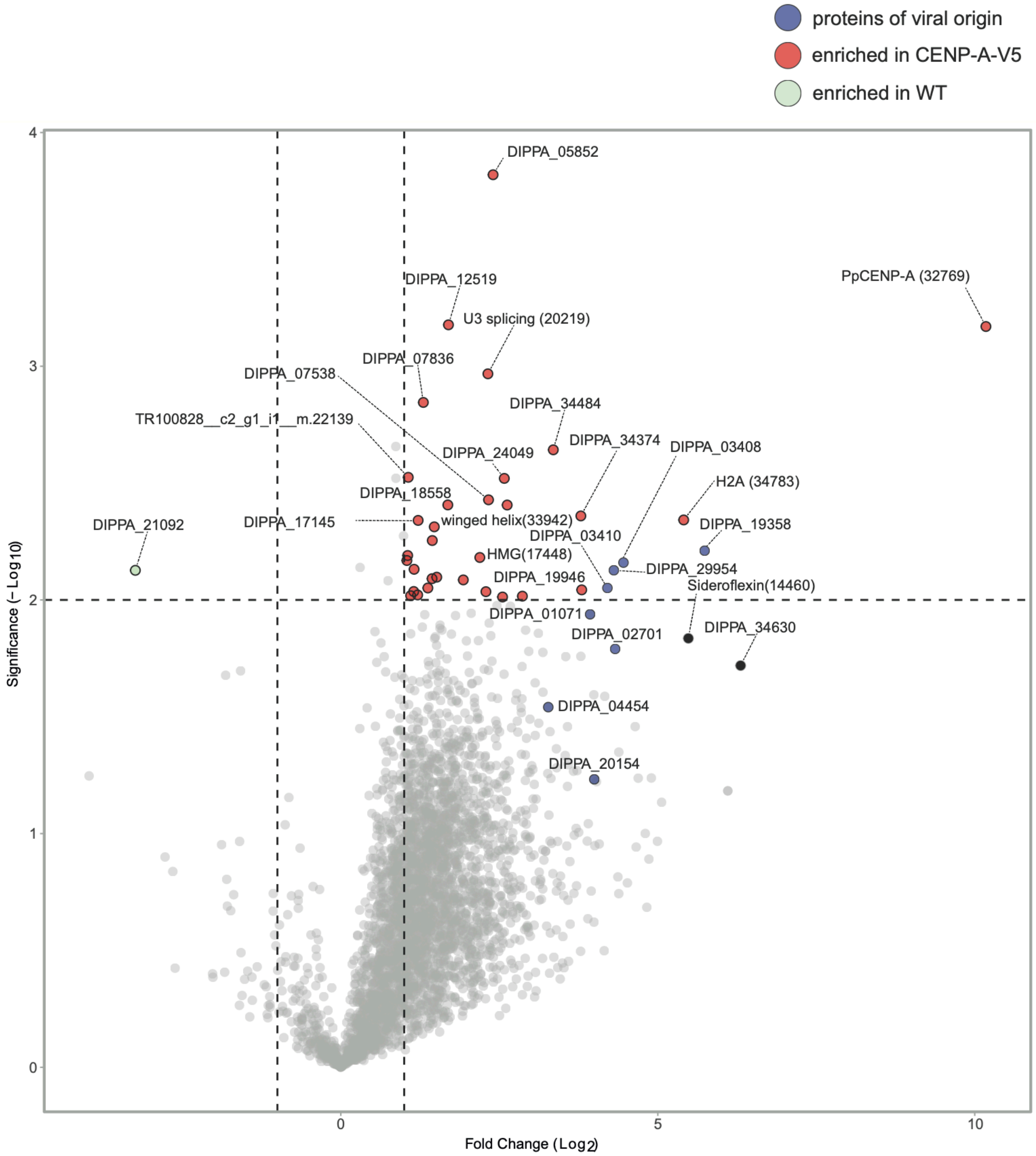


**Fig. S7 Volcano plot of CENP-A-V5 pulldown.** Enrichment analysis of V5 pulldown on WT vs CENP-A-V5 cells. Experiment was performed in triplicate. Dotted lines represent a significance threshold of 0.01 and a fold change threshold of 1.5. No strong interactors were found beyond CENP-A itself. Homologs of prophage tail fiber subunits are indicated in purple, showing modest enrichment in CENP-A vs WT.

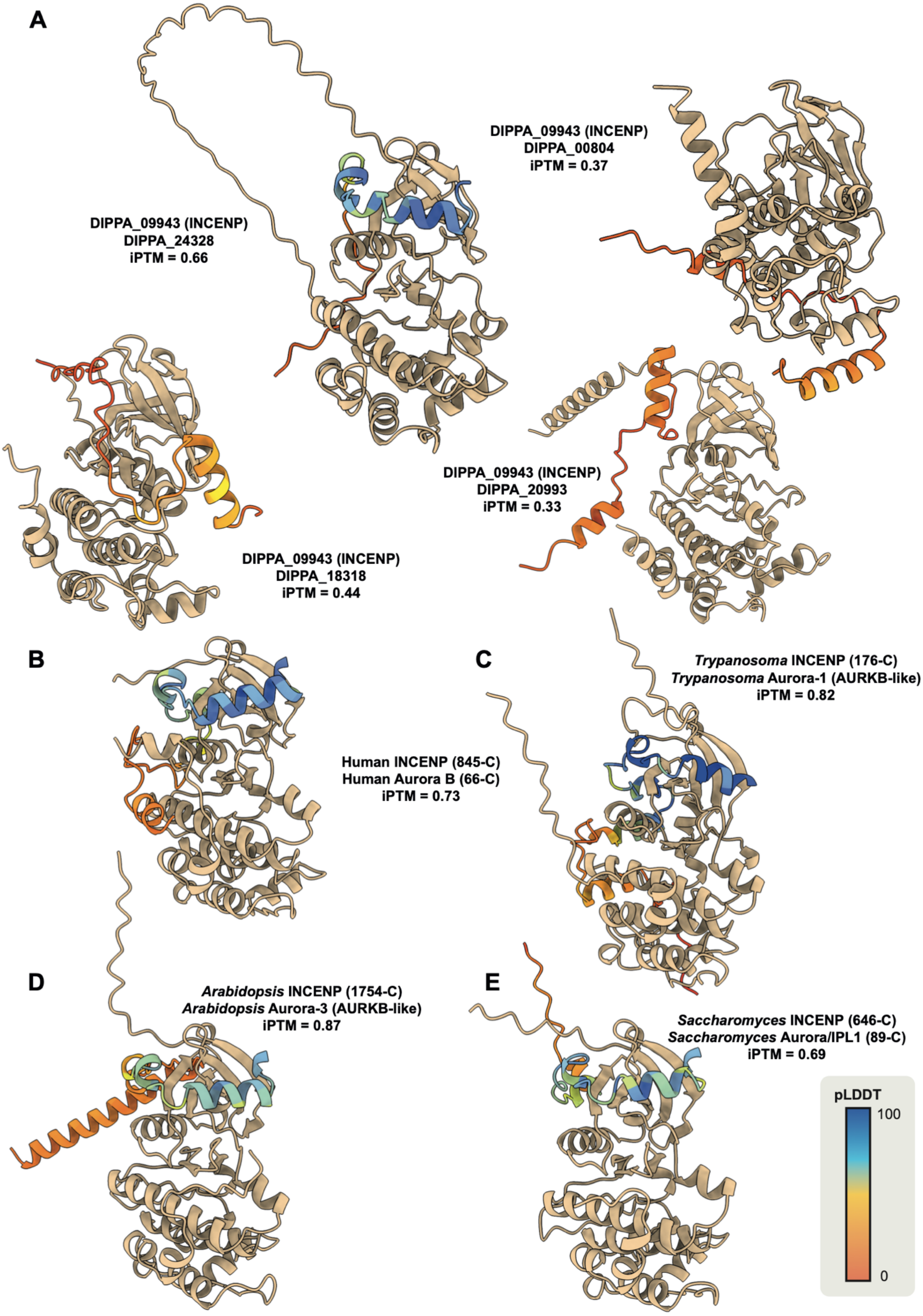


**Fig. S8 INCENP-Aurora interactions as modelled by AlphaFold3.** For model eukaryotes (human, yeast, plant, trypanosoma) and *P. papillatum,* we predicted the interaction of INCENP with an Aurora kinase. For the model eukaryotes in which the designated Aurora kinase (Aurora B) is known, high iPTM scores (>0.6) were obtained. For *P. papillatum,* the highest iPTM score was found for DIPPA_24328, which according to the phylogram in Fig. S6 is supposedly the paralog of Aurora-3/AUK3 in *T. brucei*. Note the same orientation of DIPPA_24328 and INCENP relative to each other compared to those of Aurora B and INCENP in the model eukaryotes. Aurora kinases are depicted in the colour 'sand', while INCENP shows the strength of the AlphaFold prediction (pLDDT score - as per 4 bins of the AF3 webserver).

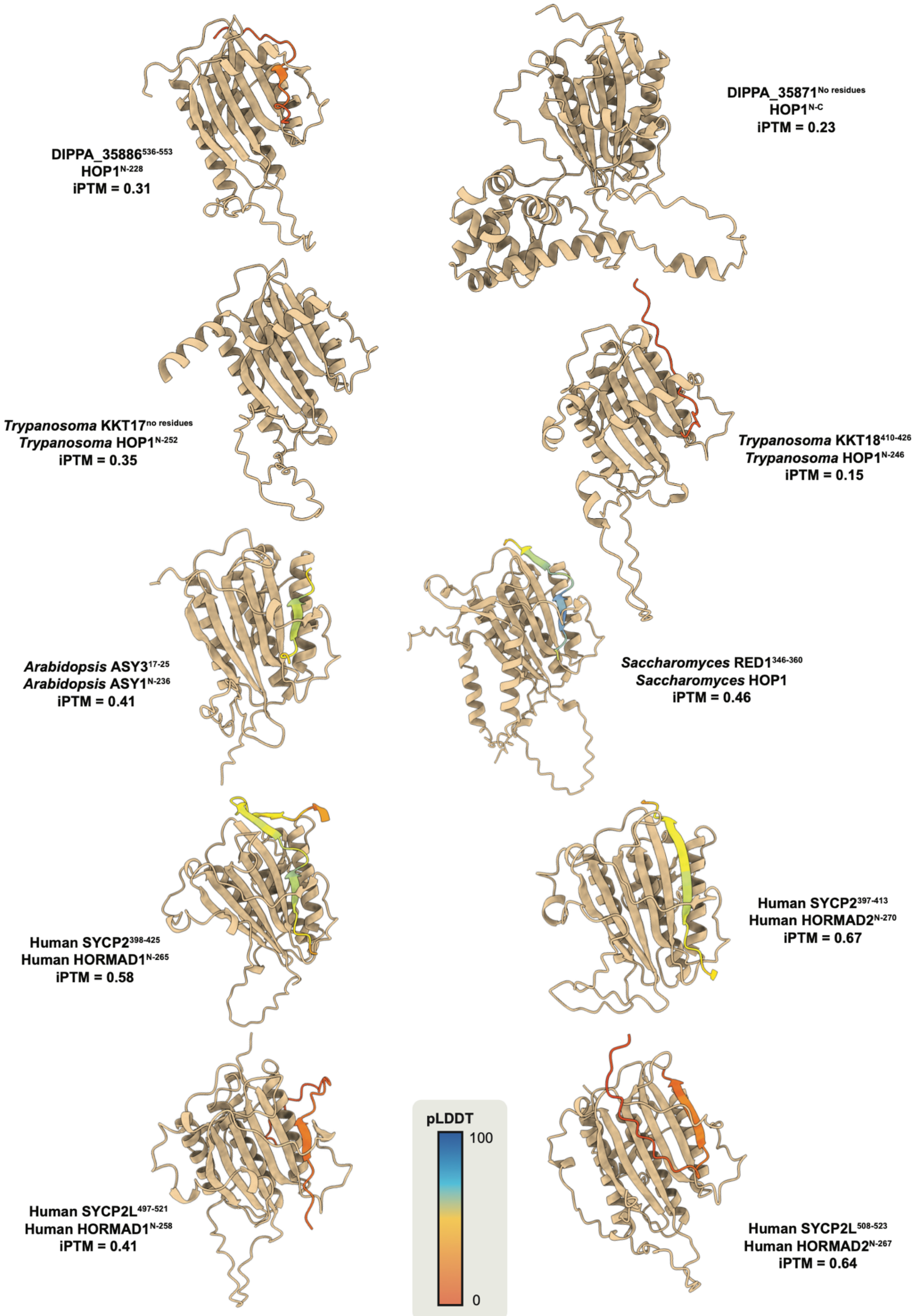
DIPPA_35886$^{536-553}$
HOP1$^{N-228}$
iPTM = 0.31
DIPPA_35871$^{No\ residues}$
HOP1$^{N-C}$
iPTM = 0.23
*Trypanosoma* KKT17$^{no\ residues}$
*Trypanosoma* HOP1$^{N-252}$
iPTM = 0.35
*Trypanosoma* KKT18$^{410-426}$
*Trypanosoma* HOP1$^{N-246}$
iPTM = 0.15
*Arabidopsis* ASY3$^{17-25}$
*Arabidopsis* ASY1$^{N-236}$
iPTM = 0.41
*Saccharomyces* RED1$^{346-360}$
*Saccharomyces* HOP1
iPTM = 0.46
Human SYCP2$^{398-425}$
Human HORMAD1$^{N-265}$
iPTM = 0.58
Human SYCP2$^{397-413}$
Human HORMAD2$^{N-270}$
iPTM = 0.67
Human SYCP2L$^{497-521}$
Human HORMAD1$^{N-258}$
iPTM = 0.41
pLDDT
100
0
Human SYCP2L$^{508-523}$
Human HORMAD2$^{N-267}$
iPTM = 0.64

**Fig. S9 SYCP2-Hop1 interactions as modelled by AlphaFold3.** For model eukaryotes (human, yeast, plant, trypanosoma) and *P. papillatum,* we predicted the interaction of SYCP2 with the meiotic protein Hop1. For the model eukaryotes in which the designated SYCP2 and Hop1 orthologs are known, iPTM scores are relatively higher (>0.4). For *P. papillatum,* the highest iPTM score was found for DIPPA_35886 (SYCP2L2; 0.31). This score is low (0.6<iPTM<1: good score), indicating that an interaction with Hop1 might be present for DIPPA_35886, but is not certain. For SYCP2L1, no interaction was predicted. Hop1 orthologs are depicted in the colour 'sand', while the closure motif of the SYCP2 orthologs shows the strength of the AlphaFold3 prediction (pLDDT score - as per 4 bins of the AF3 webserver).

### TABLES

**Table S1:** source of predicted proteins of eukaryotic transcriptomes and genomes of various eukaryotes used in this study

**Table S2:** Paradiplonema ID infosheet

**Table S3:** HHsearch searches

a: initial search (KT/KKT vs. Diplonemea OGs)

reciprocal search (top 10 Diplonemea OGs ←→ Diplonemea OGs + KOG/COG + PFAM + KT/KKT)

c: bidirectional hits from reciprocal search

**Table S4:** HMMsearch –max

**Table S5:** Foldseek searches

a: initial search (KT/KKT from representative species vs. *P. papillatum* AF2 structures).

b: reciprocal search (top 10 *P. papillatum,* AF2 structures ← → KT/KKT from representative species + full proteomes of *A. thaliana, H. sapiens, S. cerevisiae* and *T. brucei*).

c: bidirectional hits from reciprocal search

**Table S6:** Discoba kinetochore evolution - presence/absence table

**Table S7:** Pulldown of CENP-A-V5

**Table S8:** Primer sequences used for endogenous tagging

### FILES

**File S1:** cytoscape network file for HHsearch

**File S2:** cytoscape network file for Foldseek

**File S3:** Figshare repository: https://figshare.com/s/a3c0f6af159c4545f393 containing,

- Complete set of predicted structures for all *P. papillatum* folds predicted with AF2
    - *P_papillatum_AF2_foldome_rank1_pdbs.tar.gz*
- HH-suite databases used in the homology search bioinformatics pipelines of KKT alignments, cKT alignments and diplonemid orthologous groups
    - *kkt_kkip_hhsearch_db.tar.gz*
    - *ckt_hhsearch_db.tar.gz*
    - *Diplonemea_hhsearch_db.tar.gz*
- Foldseek database of the full *P. papillatum* foldome
    - P_papillatum_af2_foldseek_db.tar.gz
    - the full foldomes of the representative species + the *P. papillatum* foldome + the curated kinetochore folds: *repspecies_ppal_kkt-kt_foldseek_db.tar.gz*
- Curated set of AlphaFold2-predicted structures of kinetochore proteins
    - *kkt-kt_foldseek_db.tar.gz*
- Output files from hhsearch and foldseek as performed during the homology searches
    - *hhsearch_output.tar.gz*
    - *foldseek_output.tar.gz*
- Transcriptomes of Diplonemea spp. used in the bioinformatics pipeline
    - *Diplonemea_sequence_database.tar.gz*
- OrthoFinder output run on the Diplonemea dataset
    - *orthofinder_diplonemea.tar.gz*

**File S4:** Zip file for CENP-A-V5 confocal images
