## Supplementary figures and images for "On the possibility of yet a third kinetochore system in the protist phylum Euglenozoa"

### FileS4_CENP-A-V5_1.tif

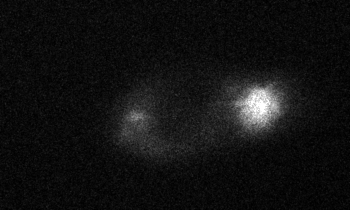

### FileS4_CENP-A-V5_2.tif

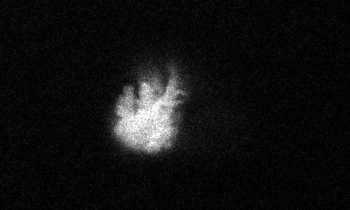

### FileS4_CENP-A-V5_3.tif

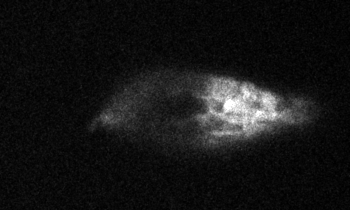

### FileS4_CENP-A-V5_4.tif

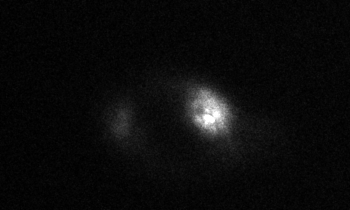
